# The transposition rate has little influence on equilibrium copy numbers of the P-element

**DOI:** 10.1101/2021.09.20.461050

**Authors:** Robert Kofler, Viola Nolte, Christian Schlötterer

**Affiliations:** Institut für Populationsgenetik, Vetmeduni Vienna, Veterinärplatz 1, 1210 Wien, Austria

## Abstract

The invasion of transposable elements (TEs) in mammals and invertebrates, is likely stopped by piRNAs that emerge after insertion of the TE into a piRNA cluster (the trap model). It remains, however, still unclear which factors influence the dynamics of TE invasions. The activity of the TE (i.e. transposition rate) is one of the frequently discussed key factors. Here we take advantage of the temperature-dependent activity of the P-element, a widely studied eukaryotic TE, to test how TE activity affects the dynamics of a TE invasion. Taking advantage of the temperature-specific activity of the P-element, we monitored its invasion dynamics in experimental *Drosophila simulans* populations at hot and cold culture conditions. Despite marked differences in transposition rates, the P-element reached very similar copy numbers at both temperatures. The reduction of the insertion rate upon approaching the copy number plateau was accompanied by the emergence of similar amounts of piRNAs against the P-element at both temperatures. Interestingly, the ping-pong cycle, which degrades TE transcripts, becomes only active after the copy number has reached the plateau. We show that the P-element abundance is an order of magnitude lower and fewer P-element insertions in piRNA clusters were observed than expected under the trap model. We conclude that the transposition rate has at the most only a minor influence on TE abundance, but other factors, such as paramutations or selection against TE insertions are shaping the TE composition of organism.

## Introduction

Transposable elements (TEs) are short stretches of DNA that spread within genomes, even when they reduce the fitness of the host [36, 22, 66]. TEs are found in most prokaryotic and eukaryotic genomes ([8, 77] and TEs frequently invade new species by horizontal transfer [55, 24, 69]. After horizontal transfer a TE may spread in a naive species until all populations are infected. The increase in copy number of the TEs is counteracted by small RNAs and negative selection. TEs may be deleterious to hosts and thus negatively selected due to three mechanisms: 1) TE insertions could directly disrupt genes or promoter regions, 2) ectopic recombination between insertions at different sites could lead to deleterious genomic rearrangements, and 3) the products of TEs such as the transposase could be deleterious (e.g., by generating DNA damage as found during hybrid dysgenesis [63, 61]).

In mammals and invertebrates the defence against TEs is mediated by piRNAs, small RNAs ranging in size between 23 to 29*nt* [12, 34, 20, 67]. These piRNAs interact with PIWI clade proteins and silence the TEs at the transcriptional and the post-transcriptional level [12, 34, 73, 52]. piRNAs are largely derived from discrete genomic loci the so called piRNA clusters [12, 30]. piRNA clusters are usually found in the heterochromatin, close to the euchromatin boundary, and may constitute substantial portions of genomes [12]. In *D. melanogaster*, for example, piRNA clusters account for about 3.5% of the genome [12]. The trap model proposes that an invading TE multiplies until one copy jumps into a piRNA cluster, which triggers the production of piRNAs that silence invading TEs [7, 57, 83, 33, 80, 23, 67].

Apart from negative selection and small RNAs, which counteract the spread of TEs, other factors may impact the dynamics of TE invasions. One frequently discussed factor is the transposition rate, i.e. the probability that a TE insertions generates a novel copy [18, 63, 44, 47]. The transposition rate varies substantially among TE families and it was long thought as the major factor determining the abundance of TEs in organisms [63, 18]. The impact of the transposition rate depends on the mechanism by which TE invasions are controlled. If TE invasions are solely counteracted by negative selection an equilibrium may result where the number of TE insertions gained by transposition equals the number of insertions lost by negative selection [18, 19, 5]. The TE abundance at this equilibrium depends on the transposition rate. In case an active TE is immediately silenced by a single insertion in a piRNA cluster, the transposition rate may only have a minor influence on the level at which TE copy numbers stabilize (i.e. the plateau level) [47, 43].

Although these two models generate clear predictions about the role of the transposition rate, several lines of evidence suggest a more complex scenario. For example, a lag-time (e.g. 7 generations) between the emergence of a cluster insertion and the establishment of an effective piRNA based defence [39] will cause an effect of transposition rate, even without selection. If both negative selection and cluster insertions control TE invasions, complex 3-way interactions among negative selection, transposition and piRNA clusters may arise where the transposition rate influences the abundance of TEs at the equilibrium level [47]. Finally it is feasible that the trap model does not hold. For example cluster insertions are not required for maintaining silencing of TEs [30] and in koalas the initial piRNAs against an invading TE arise even without cluster insertion [81]. The role of the transposition rate in alternative models of TE silencing are yet unclear.

Despite this potentially important role of the transposition rate, empirical evidence has been difficult to obtain. Comparing invasions of different TE families with varying transposition rates will only yield inconclusive results, as the families will also differ in many other factors that influence invasion dynamics such as insertion biases (e.g. into safe havens, genes, promoters), the mutation rate (including internal deletions) and the effect on the host [75, 74, 29, 27, 14].

Here we propose an elegant solution to overcome these limitations. Since the activity of the P-element, one of the most widely studied DNA transposons, varies with temperature [61, 51], we studied the P-element invasion dynamics at two different temperatures (hot and cold conditions) in experimental *D. simulans* populations. The advantage of our experimental design is that we were able to study the influence of the transposition rate on invasion dynamics while minimizing the impact of other factors. Our analyses confirm substantially different transposition rates at the two temperatures, but the abundance of the P-element plateaus at an almost identical level. This plateauing of the P-element was accompanied by a rapid emergence of piRNAs against the P-element at both temperatures. All parameters characterizing the P-element invasion, such as insertion biases, the size of piRNA clusters, the fraction of insertions with internal deletions that could down-regulate the P-element, where quite similar among the two temperatures. Modelling the invasions in our populations we show that a trap model with negative selection against TEs largely captures the observed invasion dynamics.

## Results

### Invasion dynamics at cold conditions

To investigate the role of the transposition rate during TE invasions we monitored the spread of the P-element in experimentally evolving *D. simulans* populations at hot (18 – 28°C) and cold conditions (10 – 20°C). Since the P-element activity depends on temperature [51, 61] this setup allows us to investigate the influence of the transposition rate on invasion dynamics while minimizing the influence of other TE family specific factors such as differences in transcription factor binding sites or insertion preferences. The experimental populations were established from 202 isofemale lines collected 2010 in Florida. The population from Florida was at an early stage of the P-element invasion where about 25 – 44% of the isofemale lines carried P-element insertions [49, 37]. We used 3 replicates, a census population size of 1250 and non-overlapping generations. Our previous analysis of this P-element invasion only covered 40 generations of the cold conditions, which was not sufficient to observe the plateauing of P-element copy numbers. Here, we significantly extended the experiment by additional 60 generations, which corresponds to a total of 10 years of experimental evolution. We monitored several key parameters of the cold invasion including the degree of gonadal dysgenesis, the abundance of P-element insertions and the amount of piRNAs against the P-element. All data, previously published ones and newly generated ones, were (re)analyzed using the same pipeline to ensure comparability of the data.

We first asked whether the P-element reached stable copy numbers in the cold invasion by generation 100. The experimental populations were sequenced in 10 generation intervals as pools (Pool-Seq [72]; for an overview of all genomic data used in this work see supplementary table S1). The number of P-element insertions per haploid genome was estimated with DeviaTE, which normalizes the coverage of the P-element to the coverage of single-copy genes [76]. The cold invasion reached a maximum of 14 – 22 P-element insertions per haploid genome around generations 60 – 70 (fig. 1A). At later generations P-element copy numbers decreased slightly to about 12 – 18 copies per haploid genome (fig. 1A; supplementary table S2; supplementary figs. S1, S2, S3). By contrast, the hot invasion reached stable copy numbers at generation 20 with about 13 – 17 copies per haploid genome (fig. 1A; supplementary table S2; supplementary figs. S4, S5, S6). The copy numbers at the plateau were very similar between hot and cold conditions (Welch Two Sample t-test, hot generation 60 vs. cold generations 100, *p* = 0.67). We measured the effective transposition rates (*u*′ = *u − x*) during the invasion since we are are not able to determine transposition rate (*u*) and selection (*x*) directly. The effective transposition rate differed among the hot and cold invasion (fig. 1B; supplementary table S2). The hot invasion is characterized by a high transposition rate for the first 20 generations (*u*′ ≈ 0.099 – 0.209) followed by a drop to about zero at later generations fig. 1B; supplementary table S2). In the cold invasion the transposition rate is more variable. A low rate for the first 30 generations (*u*′ ≈ 0.027 – 0.087) is followed by the maximum rate between 30-40 generations (*u*′ ≈ 0.091 – 0.096). Between 40-60 generations the rate is low again (*u*′ ≈ 0.017 – 0.050) and drops to values around zero after 60 generations 1B; supplementary table S2). At the early stages of the invasions, the effective transposition rates are significantly higher at hot than at cold conditions (fig. 1B; supplementary table S2; Welch Two Sample t-test, generations 0 – 10, *p* = 0.0058). To obtain estimates of the TE abundance independent of DeviaTE, we also computed the normalized number of reads mapping to each TE (rpm; reads per million). Using this alternative approach we obtained very similar invasion dynamics (supplementary fig. S7; supplementary table S2) and again found that the transposition rates are significantly higher at hot than at cold conditions (Welch Two Sample t-test, generations 0 – 10, *p* = 0.0074) whereas the final abundance of the P-element is similar between the hot and the cold invasions (Welch Two Sample t-test: hot generation 60 vs. cold generation 100, *p* = 0.96).

**Figure 1:**
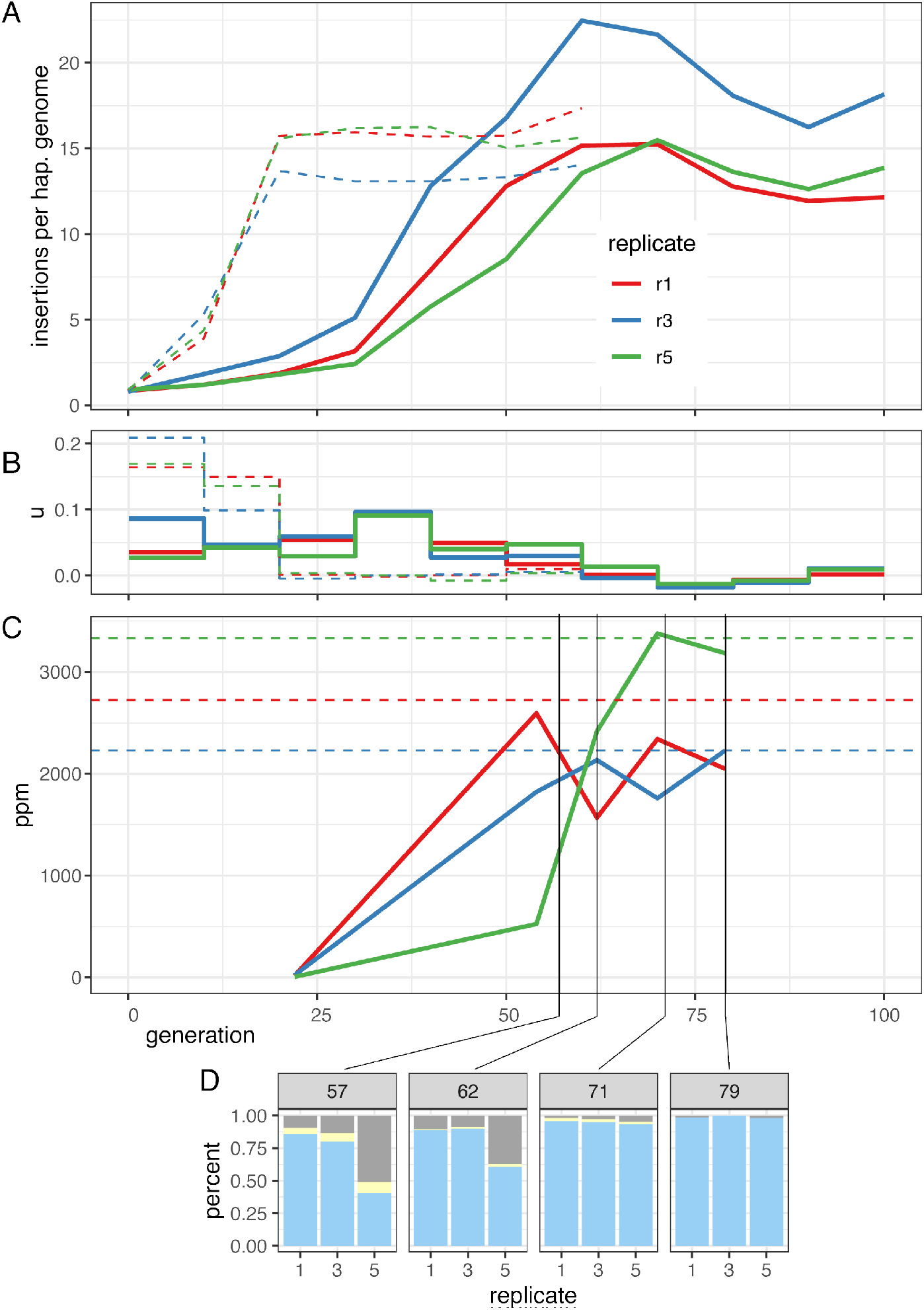
Dynamics of a natural P-element invasion in *D. simulans* at cold conditions (solid lines). A) Abundance of P-element insertions during the invasion in three replicates. Hatched line shows the previously published invasion dynamics at hot conditions [51]. B) Effective transposition rates (*u*′) during the invasions. C) Abundance of piRNAs complementary to the P-element (ppm piRNAs per million miRNAs). Hatched lines show the average abundance of P-element piRNAs at the plateau (≥ 20 generations) of the hot invasion. Note that similar levels of piRNAs are necessary to stop the invasions at hot and cold conditions. D) Abundance of dysgenic ovaries at 29°C. blue: normal ovaries, grey: dysgenic ovaries, yellow: intermediate

We thus conclude that the P-element reached similar copy numbers at the hot and the cold invasions despite transposition rates being significantly lower in cold than in hot conditions.

### piRNAs

According to the trap model the invasion of a TE is stopped when one copy of the TE inserts into a piRNA cluster, which triggers the production of piRNAs complementary to the TE. This prediction can be scrutinized by monitoring the emergence of piRNAs complementary to the P-element during the invasion. We analysed small RNAs from whole bodies of female flies at generations 22, 54, 62, 70, 79 of the cold invasions and generations 22, 44, and 108 of the hot invasions (data from generations 62, 70 and 79 of the cold invasion were generated in this work; for an overview of all small RNA data used in this work see supplementary table S3). In the cold invasions the number of piRNAs complementary to the P-element rapidly increased between generations 22 and 62 (fig. 1C; supplementary table S3), consistent with the decrease of the transposition rate after generation 60. Similarly to the hot invasion [51] only the abundance and piRNA level of a single TE, i.e. the P-element, increased during the cold invasion (supplementary fig. S13). The piRNAs were mostly anti-sense to the P-element with a length between 23 and 29nt (supplementary fig. S8) and distributed over the length of the P-element (supplementary fig. S9).

The piRNA response is amplified by the ping-pong cycle [12, 34], and we evaluated to what extent this process was driving the increase in piRNA copies during our experiment. The ping-pong cycle is based on two PIWI clade proteins Aub and Ago3. During the ping-pong cycle, RNA cleavage products of Aub are loaded onto Ago3 and vice versa. Since the Aub cleavage site is shifted by 10bp from the Ago3 cleavage site a characteristic peak at position 10 can be found when plotting the average distance between the 5’ ends of sense and antisense piRNAs (i.e. the ping-pong signature; [12, 34]). The size of the peak indicates the contribution of the ping-pong pathway to the total piRNA population. We found a peak at position 10 and a peak at position 12 (supplementary fig. S10). This peak at position 12 is mostly due to highly abundant piRNAs at positions 1162 and 1164 of the P-element. When excluding these piRNAs a ping-pong signal at position 10 can be observed for cold evolved populations at generations ≥ 54 (supplementary fig. S11). The amount of piRNAs in the cold invasion at generation 22 was too small for computing ping-pong signatures. In both the hot and the cold invasion the ping-pong signature was weakest in the earliest generation for which the signature could be computed (hot 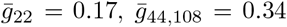, Wilcoxon rank sum test *W* = 1, *p* = 0.047; cold 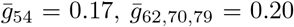, Wilcoxon rank sum test *W* = 9, *p* = 0.48; supplementary figs. S11, S12). We observed clear ping-pong signatures only after the P-element copy number stabilized. This suggests that secondary piRNAs are not the main cause for the reduced transposition rate at the plateauing phase. We propose that the ping-pong cycle may be more important for robust silencing of the P-element at later generations.

The amount of piRNAs at the plateau of the cold invasions is slightly lower, than the level of piRNAs at the plateau of the hot invasions (fig 1C; average piRNAs in ppm, hot generations 22-108 *r*1 = 2723, *r*3 = 2229, *r*5 = 3331; cold generations 70-79 *r*1 = 2193, *r*3 = 1994, *r*5 = 3279). This is surprising given that the P-element expression is higher at hot than at cold conditions [51] (copy numbers at the plateau are similar at both conditions). This suggests that either more piRNAs are produced than required for silencing or that piRNAs silence more efficiently at hot conditions.

Even when a TE is silenced by the piRNA pathway and the copy number remains stable, it remains an open question whether the TE is silenced in all individuals of a population or if the TE remains active in a small subset in the population. Our inter-population assays of gonadal dysgenesis, which estimate the fraction of females with atrophied ovaries, suggest that silencing of the P-element may be heterogenous within the populations. Gonadal dysgenesis (GD) estimates the fraction of females with atrophied ovaries. GD is expected for females which carry functional P-elements but produce insufficient amounts of piRNA to repress the P-element [13, 42]. At generations 57 and 62 between 9-17% of the ovaries are dysgenic in replicates 1 and 3 and at generation 79 the total number of dysgenic ovaries in these two replicates drops to 0.0-1.2%. This confirms that the cold invasion is largely silenced at generation 70. Interestingly, the difference in atrophied ovaries between generation 62 and 79 is only to a minor extent reflected by the levels of piRNA (fig. 1C,D), but by an increase of the ping-pong signature (supplementary fig. S11). Since we measured piRNAs levels from pooled flies, but estimated dysgenesis in individual females, our results suggest that the distribution of piRNAs differs among individual females. This may reflect the segregation of different cluster insertions in the evolving populations. At later generations, the cluster insertions may be more homogeneously distributed among individual flies, resulting in more females without hybrid dysgenesis. An alternative interpretation is that the production of the initial piRNAs against an invading TE is instable with stochastic fluctuations in piRNA levels. Only when the ping-pong cycle is fully established, a robust piRNA silencing is achieved across all females.

If the trap model holds, P-element insertions in piRNA clusters should emerge during the invasion, especially at the plateau of the invasions [47]. Therefore we determined the positions of piRNA clusters in a long-read assembly of *D. simulans* [17]. We aligned all small-RNA data from the cold invasion to the genome and identified regions with a high density of piRNAs (i.e. piRNA clusters) using a previously described algorithm that maximizes a local-score [51, 26]. The piRNA clusters in *D. simulans* have a total size of 4.24*Mb* which corresponds to about 2.6 % of the genome (assuming a genome size of 162*Mb* for *D. simulans* [11]). The positions of P-element insertions were inferred from the Pool-Seq data using PoPoolationTE2 [50]. The first P-element insertions in piRNA cluster were observed in all replicates at generation 50 of the cold invasion (supplementary fig. S14, S15, S16). Next we asked if the observed number of cluster insertions is sufficient to stop an invasion under the trap model. Even under the assumption that one cluster insertion per diploid individual is biochemically sufficient to silence a TE, the number of cluster insertions necessary to stop an invasion in a population will be higher when cluster insertions are segregating as predicted by the trap model [47, 43]. Segregation of cluster insertions leads to an uneven distribution of the insertions in a population. As a consequence some individuals in a population will have many cluster insertions while others will end up without any cluster insertion and the TE will be active in these individuals [47]. When diploid individuals carry on the average four cluster insertions, the vast majority of the individuals within a population will end up with at least one cluster insertion (hence the TE is largely silenced throughout the population). When one cluster insertion is fixed, then two insertions per diploid individual are sufficient to silence a TE [47]. Since all cluster insertions in the cold and the hot invasion are segregating at a low population frequency (supplementary fig. S14, S15, S16, S17, S18, S19), two cluster insertions per haploid genome will be necessary to stop the invasion in our experimental populations.

Using stringent criteria (unambiguously mapped piRNAs to define the location of piRNA clusters and unambiguously mapped reads to locate TE insertions) the mean number of cluster insertions per haploid genome was 0.21 at the plateau of the cold invasion (generations 70 – 100) and 0.24 at the plateau of the hot invasion (generation 20 – 60; supplementary table S5), which is insufficient to stop an invasion under the trap model. One possible explanation for the low number of cluster insertions may come from telomere associated sequences (TAS) [84]. TAS are located at the ends of the major chromosome arms in Drosophila and likely act as piRNA clusters [3]. Since TAS largely consist of repetitive sequences, it is difficult to identify TE insertions based on unambiguously aligned reads in these regions [84]. To find TE insertions in TAS we first extracted the regions between the most distal gene and the end of each chromosome arm, yielding in total five sequences of TAS regions (X-TAS, 2L-TAS, 2R-TAS, 3L-TAS, 3R-TAS; for coordinates see supplementary table S6) From our paired-end reads we additionally extracted all reads where only one mate maps to the P-element (but not the read itself). These reads enabled us to anchor P-element insertions in a reference sequence (henceforth “anchor reads”). Alignment of these anchor reads to the five TAS sequences allowing for ambiguously mapped reads showed that P-element insertions are indeed highly enriched in TAS (5.9-fold at the plateau of the cold invasion; supplementary table S7). This enrichment was most pronounced in X-TAS followed by 3R-TAS (*X* = 16.7, 3*R* = 10.1; supplementary table S7). A strong enrichment of P-element insertions in X-TAS was also previously observed [40, 84, 1]. At the plateau the enrichment of P-element insertions in TAS was more pronounced at hot than at cold conditions (hot: *all* = 7.3, *X* = 41.7, 3*R* = 19.2; cold: *all* = 5.9, *X* = 16.7, 3*R* = 10.1; Wilcoxon rank sum test for all three sets *p* < 0.01; supplementary table S7). Nevertheless, even when TAS insertions are considered, the average number of insertions per haploid genome is still not sufficient for stopping the P-element invasions under the trap model (cold =0.37, hot =0.64; not counting TAS insertions twice; supplementary table S5). Since we cannot rule out that cluster insertions in highly repetitive regions were missed, we estimated the number of cluster insertions using the most liberal criteria: we considered ambiguously mapped small RNAs for defining the positions of piRNA clusters and considered ambiguously mapped anchor reads within TAS as well as piRNA clusters. Even with these liberal criteria each individual carries only about 1.46 cluster/TAS insertions at cold conditions and 1.63 at hot conditions, still less than the two insertions required for reliable silencing of the P-element.

Although we used a long-read based assembly, some piRNA clusters may be missing in our assembly. Furthermore piRNA clusters are likely polymorphic [79, 78] and therefore the sequences of piRNA clusters in our experimental populations may differ from the assembled ones. At cold (hot) conditions about 8.1% (11.2%) of the anchor reads can not be aligned to the reference genome (supplementary table S9). Assuming that equal proportions of aligned and not-aligned anchor reads map to clusters/TAS then each individual could, as an upper estimate, carry about 1.58 cluster/TAS insertions at cold and 1.81 at hot conditions (at the plateau). The actual number of cluster insertions may only be higher than this estimate if i) a large fraction of the not-aligned anchor reads maps to piRNA clusters or ii) we missed some piRNA clusters, e.g. strain specific clusters or lowly-expressed clusters.

To summarize we estimate that at the plateau of the invasion each individual carries between 0.21 (stringent criteria) and 1.8 (most liberal criteria) cluster insertions per haploid genome, which is smaller than the 2 insertions expected under the trap model. Although piRNAs against the P-element and P-element insertions in piRNA clusters rapidly emerged in our experimental populations the number of cluster insertions is probably not sufficient to control an invasion under the trap model.

### Differences among hot and cold conditions

We showed that the transposition rate had little influence on P-element copy numbers at the plateau of an invasion. Since differences in transposition rate were modulated by temperature it is important to test whether temperature affects other factors influencing the invasion dynamics of the P-element.

We first investigated the influence of internal deletions (IDs) of the P-element since IDs may repress P-element activity and our previous work showed that IDs emerge faster at hot than at cold conditions [51]. IDs, such as KP or D50 [9, 71] produce non-functional transposases that retain the DNA-binding capacity but are unable to mobilize the P-element. KP-like repressors prevent functional transposases from mobilizing the P-element by blocking transposase binding sites (supplementary fig. S20A, green shade in lower panel). We measured the dynamics of IDs in hot and cold condition using DeviaTE allowing for 3bp tolerance in the position of IDs (the exact position of ID is frequently ambiguous in alignments). We found 65 distinct IDs in the cold invasion and 61 in the hot invasion. Nine IDs shared the same break points in multiple replicates. Assuming that identical break points rarely arise independently, we conclude that these IDs were already present in the base population (7 IDs were detected in the cold invasion; supplementary fig. S20A, dashed lines).

Some IDs increased in frequency within the P-element population (supplementary fig. S20B). This frequency increase may be driven either by positive selection resulting from the repression of the P-element activity or by preferential mobilization compared to other P-elements. Especially, the ID at position 187 – 1967nt was rapidly rising in frequency in two replicates, accounting for up to 30 – 40% of all P-element copies at the last generation (supplementary fig. S20B, red). This ID is probably no KP-like repressor as its DNA binding domain is largely deleted (supplementary fig. S20, red; orange shade in lower panel A is the DNA binding domain). This makes positive selection unlikely. Rather, preferential mobilization of this ID is a more plausible explanation for the copy number increase of this ID.

Although IDs are emerging faster at hot than at cold conditions [51], the final abundance of IDs and KP-like repressors is similar at the two temperature conditions (fig. 2A; Wilcoxon rank sum test; all IDs *W* = 130, *p* = 0.052; KP-like repressors *W* = 99, *p* = 0.68). It is thus also unlikely that P-elements are differently regulated by IDs in the two temperature regimes.

**Figure 2:**
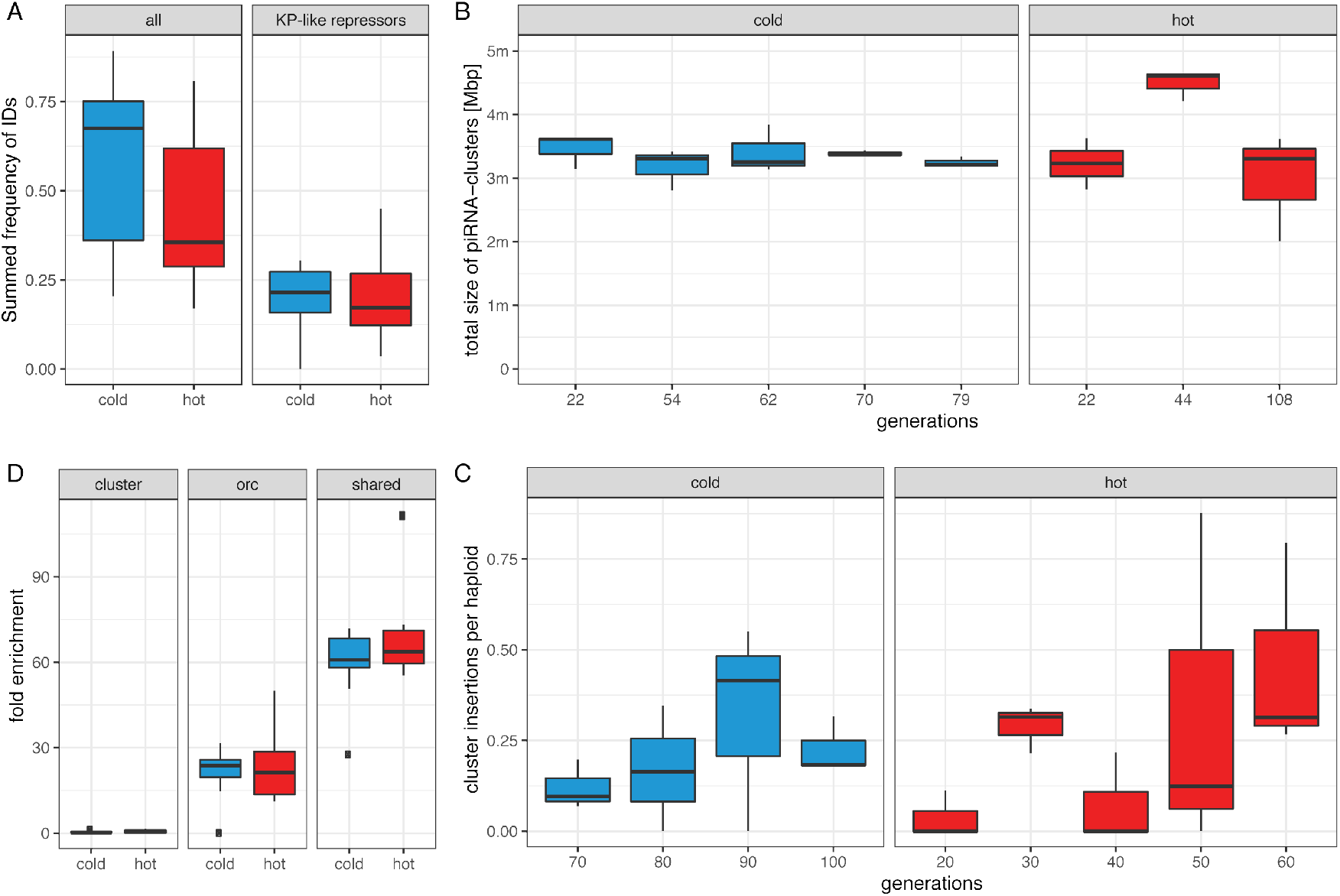
Temperature has a minor influence on some important factors governing the invasion dynamics. A) Fraction of P-element insertions with any internal deletion (ID; all) and an ID that may repress P-element activity (KP-like repressor). Only samples at the plateau of the invasions were used (hot ≥ 20, cold ≥ 70 generations). B) Total size of piRNA clusters during the invasions based on unambiguously mapped piRNAs. C) Average number of P-element insertions in piRNA clusters per haploid genome. Only samples at the plateau of the invasion are shown. D) Enrichment of P-element insertions in piRNA clusters, ORCs and insertion sites shared among species. Only samples with more than 10 P-element insertions per haploid genome were used (generations ≥ 20 hot and ≥ 50 cold).

Simulations of TE invasions with piRNA clusters showed that the size of piRNA clusters is the major factor determining the plateauing level of TE invasions [48, 43]. As the activation of some piRNAs cluster may depend on temperature [16] we tested whether differences in the size of piRNA clusters could have influenced the invasion dynamics. We estimated the size of piRNA clusters in our small RNA data based on unambiguously mapped piRNAs. The size of the piRNA cluster is very similar among hot and cold conditions (Wilcoxon rank sum test *W* = 83.5, *p* = 0.36; fig. 2B). Additionally, the abundance of P-element insertions in piRNA clusters identified using small-RNA data (stringent conditions, with unambiguously mapped piRNAs and TE insertions) is also similar between the two temperature regimes – irrespective of whether piRNAs from the hot or cold conditions were used to identify piRNA clusters (cold piRNAs, Wilcoxon rank sum test *W* = 61, *p* = 0.44; fig. 2C; hot piRNAs, Wilcoxon rank sum test *W* = 64, *p* = 0.82).

It has been suggested that the P-element has an insertion bias into diverse genomic features such as promoters of genes, TAS and origin of replication complexes (ORC) [74, 6, 40, 49]. These insertion biases may enhance (e.g. ORC) or counteract (e.g. piRNA clusters including TAS) the propagation of the P-element. Differences in the insertion bias among hot and cold conditions could thus influence invasion dynamics.

We showed that the P-element has a stronger insertion bias into TAS at hot than at cold conditions. This does not necessarily imply that an invasion is more readily silenced at hot than at cold conditions, since any insertion in a piRNA clusters (not just in TAS) may trigger the production of piRNAs that silence the P-element. Therefore we tested whether the insertion bias into any piRNA producing loci differs between the two temperatures (including piRNA producing TAS regions; stringent conditions, with unambiguously mapped piRNAs and TE insertions). However the overall insertion bias into piRNA clusters was similar between hot and cold conditions (mean enrichment for samples with more than 10 insertions: *hot* = 0.63, *cold* = 0.33; Wilcox rank sum test *W* = 91, *p* = 0.19; fig. 2D). At both temperatures the insertion bias was < 1 implying that P-element insertions in piRNA clusters are actually less abundant than expected by chance.

The insertion bias into origin of replication complex (ORC) regions may enhance the transmission of the P-element into the next generation by enabling the P-element to be duplicated multiple times during S-phase [74]. We found a strong enrichment of P-element insertions in ORC regions at both temperatures. The extent of the enrichment was very similar between hot and cold conditions (mean enrichment for samples with more than 10 insertions: *hot* = 23.2, *cold* = 21.9; Wilcox rank sum test *W* = 136, *p* = 0.72; fig. 2D).

We previously reported that some insertion sites of the P-element are shared between *D. simulans* and *D. melanogaster* [49], suggesting a strong insertion bias in both species. It is not clear if this insertion bias influences the propagation of the P-element. In our experimental populations the P-element showed the strongest enrichment at these shared sites but the extent of the enrichment is similar between hot and cold conditions (mean enrichment for samples with more than 10 insertions: *hot* = 70.2, *cold* = 60.6; Wilcox rank sum test *W* = 98, *p* = 0.30; fig. 2D). The insertion bias thus had a similar influence on the invasion dynamics of the P-element at both temperatures.

In summary we conclude that the investigated factors, such as the size of piRNA clusters, the abundance of KP-like repressors and the insertion bias, were highly similar between the temperature regimes and cannot explain the temperature-specific invasion dynamics.

### Modelling the P-element invasion

The trap model provides a good framework to predict the invasion dynamics when the key parameters insertion rate and piRNA cluster size are known. We assumed that TEs multiply at a constant transposition rate (*u_cold_* = 0.051, *u_hot_* = 0.154; i.e. the average *u* before plateauing of the invasions; supplementary table S2) in individuals without a cluster insertion and at a rate of *u* = 0 in individuals with at least one cluster insertion. We simulated five chromosome arms of 32.4*Mb* and a piRNA cluster of size 850*kb* at the end of each chromosome arm, resulting in a total genome size of 162MB and a total size of piRNA clusters of 4.25Mb. These conditions resemble our scenario in *D. simulans* where piRNA clusters also account for 2.6% of the genome. Additionally, we used a recombination rate of 4*cM/Mb* [38], a population size of *N* = 291 [4], and triggered the invasions by randomly distributing 498 TE insertions with a population frequency of 1/2 ∗ *N* in the initial population (similarly to our base population). Simulations with neutral TE insertions (*x* = 0; hence *u*′ = *u*) capture the sharp increase in TE copy numbers but fail to predict the timing as well as the level of the plateauing (supplementary fig. S21). Both the observed onset and level of the plateauing (hot 15.1 copies, 20 generations; cold 15.2 copies, 70 generations) were substantially overestimated by the simulations (hot 160 copies, ≈ 125 – 250 generations; cold 102.7 copies, ≈ 125 – 250 generations; supplementary fig. S21). Furthermore, the number of P-element insertions per haploid genome was slightly lower in the cold than in the hot simulation.

One possibility that could account for this discrepancy is that we underestimated the size of piRNA clusters. However, additional simulations show that a plateauing at about 15 insertions per haploid genome is only expected when the piRNA clusters account for 30-40% of the genome (supplementary fig. S22). Typical estimates for the size of piRNA clusters range from 3% in Drosophila to about 0.1% in humans, mice and rats [12, 32], which suggests that a larger piRNA cluster cannot explain the discrepancy.

Given this poor fit to the data, we considered a more complex trap model, which also accounts for selection against TE insertions. To avoid an equilibrium state between transposition, selection and piRNA clusters (TSC-balance) we assumed that cluster insertions are neutral [47]. In the presence of purifying selection, we cannot observe the transposition rate directly, but rather the effective transposition rate (*u*′ = *u − x*) which is the net result of transposition (*u*) and negative selection against TEs (*x*). The same effective transposition rate may be achieved by many different combinations of transposition rates and negative effects against TEs. Therefore we performed simulations of TE invasion with different combinations of these two factors. The strength of negative selection had indeed a pronounced influence on the invasion dynamics, even when the effective transposition rate was kept constant (supplementary fig. S23). A reasonable fit between expected and observed data was obtained when the negative effect of TE insertions was about *x* = 0.02, i.e. when a single TE insertion reduced host fitness on the average by 2% (supplementary fig. S23). Under this model the onset and level of the plateauing (hot 19.1 copies, ≈ 30 – 40 generations; cold 11.3 copies, ≈ 80 – 90 generations) approach the observed ones (15.1 copies, 20 generations; cold 15.2 copies, 70 generations; fig. 3). Nevertheless, some differences remain-the prominent overshooting in the hot environment and the slight underestimation of the plateauing copy number in the cold.

**Figure 3:**
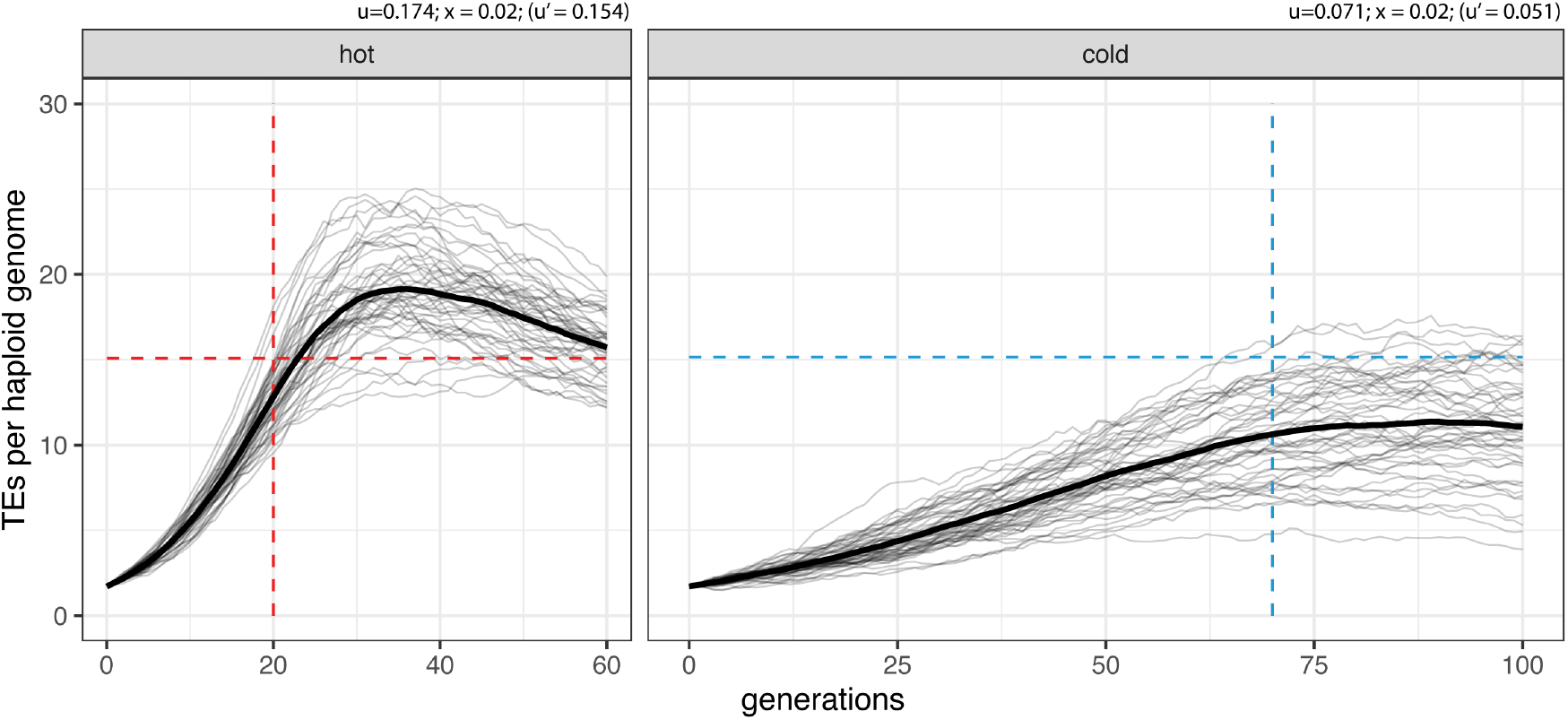
Expected and observed invasion dynamics under the trap model with negative selection against TE insertions (*x* = 0.02). The dashed lines indicate the observed onset and level of the plateau in our experimental populations. Solid lines show the expected invasion dynamics for 50 replicates and the bold lines are averages.․ *u* transposition rate, *x* negative effect of a TE insertion, *u*′ effective transposition rate.

We showed that it is feasible to generate a simple trap model with few parameters that largely captures the invasion dynamics. We caution, however, that many open questions remain about factors influencing invasion dynamics and therefore other models, for example using other combinations of parameters, may fit the data equally well.

## Discussion

We studied the influence of transposition rates on the invasion dynamics of the P-element. We show that different transposition rates, modulated by the ambient temperature of experimental *D. simulans* populations, has little influence on the number of TE insertions at the plateau. Since temperature may not only affect the transposition rates, but also other factors, we evaluated a broad range of factors (IDs, insertion bias, size of piRNA clusters) and only found a higher insertion bias into TAS regions, especially X-TAS, at hot than at cold conditions.

A preferential insertion into TAS regions at hot conditions might lead to a faster silencing of the invasion and thus a lower plateau level at hot conditions. However since the TAS regions are a subset of the piRNA producing loci and the overall insertion bias into any piRNA producing locus is similar among the two temperatures (fig. 2D), similar plateauing levels at the two temperatures are expected. In agreement with this the hot and the cold invasions plateaued at a very similar level in our experimental populations (cold 12 – 18 insertions, hot 13 – 17 insertions; fig. 1A).

It is possible that we missed additional factors that could have had an impact on the invasion dynamics. One such factor that might differ among the cold and the hot invasion is the influence of gonadal dysgenesis (GD). The extend of GD is highly temperature dependent. GD can be readily seen at high temperatures (≈ 29°C), but is rarely observed below 24°C [46, 25, 31]. It is however not clear if GD had a notable influence on the invasion dynamics in our experimental populations. While we observed GD at high temperatures (29°C) no GD was observed at the experimental conditions (i.e. GD was observed at 18-28°C for the cold invasion but not at 18-28°C for the hot invasion and 10-20°C for the cold invasion [51]). Additionally GD within populations is likely a transient phenomenon (expected during the early stages of an invasion), that can be attributed to initially low levels of piRNA. The impact of negative selection against dysgenic females on invasion dynamics is not entirely clear. GD will reduce the fitness of males having too many P-element insertions (they induce GD) as well as of females having too few P-element insertions (they are susceptible to P-element activity). Nevertheless, it is usually assumed that GD slows the spread of the P-element [43]. By exerting some selection against P-element insertions (*x*), GD could thus have had an impact on invasion dynamics at hot conditions but probably not at cold conditions, where GD is usually not observed.

Simulations of the P-element invasion showed that a slightly extended trap model with piRNA clusters accounting for about 2.6% of a genome and deleterious TE insertions (*x* = 0.02) largely capture the invasion dynamics of the P-element (fig. 3). However this model rests on several assumptions that might not hold. First we assumed that a TE invasion is stopped when the TE jumps into a piRNA cluster (i.e. the trap model). While there is ample support for this model [83, 39, 62, 23, 47] also some doubts emerged recently [30, 81]. Notably, deletion of the three largest piRNA clusters did not lead to an increase of TE activity [30]. The authors argue that instead of cluster insertions, dispersed piRNA producing TE insertions are responsible for maintaining silencing of a TEs. Such dispersed piRNA producing TE insertions may be the result of paramutations, where maternally transmitted piRNAs mediate the conversion of dispersed TE insertions into piRNA-producing loci [53, 21, 59, 35]. Notably the origin of the initial piRNAs against an invading TE is unclear under this scenario [30]. One possibility is that the production of the initial piRNAs is triggered by an insertion into a piRNA cluster. Paramutations could have a substantial effect on invasion dynamics. The emergence of some initial piRNAs against the invading TE (possibly due to a cluster insertion) may trigger a chain-reaction, where increasing numbers of euchromatic TE insertions are converted into piRNA producing loci. In the absence of negative selection against TEs, such an accelerated silencing could account for the rapid onset of the plateauing and the low abundance of the P-elements at the plateau in our experimental populations. If paramutations contribute to the silencing of P-elements, not every individual in a population needs to carry piRNA cluster insertions as euchromatic TE insertions may also produce piRNAs. Paramutations may thus also account for the low number of cluster insertions found in our experimental populations. The influence of paramutations and piRNA cluster insertions on invasion dynamics is an important open question.

Second we assumed that the emergence of piRNAs against an invading TE leads to complete silencing of a TE reducing the transposition rate to zero. It is however feasible that TEs controlled by the piRNA pathway retain some residual activity (*u* > 0) which could have a notable effect on the invasion dynamics. The slight increase in P-element abundance between generations 90 – 100 of the cold invasion suggests that the P-element may have some residual activity, despite being largely silenced by the piRNA pathway.

Third we used a very simple selection model. The assumed fitness cost a P-element insertion of 2% (*x* = 0.02) is slightly lower than the 5.5% estimated by Mackay et al. [56] for a heterozygous P-element insertion in *D. melanogaster*. We further assumed that all TE insertions have the same effect on host fitness. However, it is likely that the fitness consequence of TE insertions varies across the genome; for example insertions within important genes are more deleterious than insertions in repetitive regions. Interestingly our simulations show that the distribution of fitness effects has little influence on the onset and the level of plateauing, but a marked influence on the TE abundance after the plateauing (supplementary fig. S24). With equal fitness effects of all TE insertions, for example, our model predicts a sharp decline in TE copy numbers at later generations. While we currently lack advanced generations to test this prediction against empirical data, nearly stable copy numbers are obtained when the negative effects of the TEs follow a distribution with a mean effect of 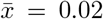 (supplementary fig. S24; equal amounts of insertions with *x* ∈ {0.0, 0.01, 0.02, 0.03, 0.04}). We assumed no epistasis among TE insertions, but it has been proposed that the fitness of individuals decreases exponentially with TE copy numbers Charlesworth and Charlesworth [18]. Such an exponential fitness decay may be mediated by the deleterious effects of ectopic recombination [19, 60]. It is an important open question whether or not ectopic recombination contributes to limiting the spread of TEs [5, 45]. Finally it is conceivable that the extend of negative selection against TE insertions differs among the temperatures. Such differences could, for example, emerge when an active transposase is generating harmful cellular effects [64] and the activity of the transposase depends on the temperature [41]. The decrease in copy numbers after the plateauing of the cold invasion, but not at the hot invasion, may be due to such temperature dependent negative effects of TEs (fig. 1A). These considerations highlight that the distribution of fitness effects of TE insertions is a crucial factor determining invasion dynamics. Unfortunately little is known about the fitness effects of TEs. Some argue that TE insertions are largely deleterious [68, 10, 56, 82], some think that most TE insertions may be neutral [2] and others argue that many TE insertions are beneficial [15]. Although many open questions remain about important parameters, modelling of TE invasions is an important step towards a quantitative understanding of TE invasions. Such a quantitative understanding might finally shed light on the question, why the TE composition varies so dramatically among species [8, 28].

## Material and Methods

### Experimental populations

Experimental populations were maintained as described previously [51]. Briefly, we collected 202 isofemale lines from a *D. simulans* population collected in November 2010 in Florida (Tallahassee). We established three replicate populations that were exposed to hot (cycling between 18°C and 28°C) and cold (cycling between 10°C and 20°C) conditions. The census population size was about 1000 and non-overlapping generations were used.

### Genomic sequencing data

We previously sequenced the hot evolved populations until generation 60 and the cold evolved populations until generation 40. All populations were sequenced as pools (Pool-Seq [72]) at each 10^*th*^ generation using the Illumina paired-end technology (Accession numbers PRJEB20533 and PRJEB20780). Here we additionally sequenced the cold evolved populations at each 10^*th*^ generation from generation 50 to 100. For each sample the DNA of pooled female and male flies was extracted using a high salt extraction protocol [58] and sheared with a Covaris S2 device (Covaris, Inc. Woburn, MA, USA). Libraries were prepared using the TruSeq DNA PCR-Free protocol (Illumina, San Diego, CA) and sequencing was performed with the Illumina HiSeq X Ten platform (Illumina, San Diego, CA). For an overview of the sequencing data used in this work see Supplementary table S1.

### small RNA

In addition to previously published small RNA data (hot at generations 22, 44, 108; cold at generations 22, 54; [51]) we sequenced small RNA from cold evolved populations at generations 62, 70, 79. As described previously we extracted total RNA from about 50 females (aged 3-4 days) reared in a common garden for 2 generations at 23°C. Sequencing of the small RNA libraries was done by Fasteris (https://www.fasteris.com/dna/). To avoid inconsistencies, all small RNA data, including the previously published ones, were analyzed using the same pipeline. We initially filtered for reads with a length between 15 and 35bp and aligned them to a database consisting of the *D. simulans* tRNAs, miRNAs, mRNAs, snRNAs, snoRNAs, rRNAs (v1.4; Flybase http://flybase.org/) as well as the consensus sequences of TEs in *D. melanogaster* (v9.42; plus Mariner GenBank:M14653.1 [70]) using novoalign (v3.03.02; http://www.novocraft.com/ -F STDFQ -o SAM -o FullNW -r RANDOM). We filtered small RNAs with more than 2 mismatches. The abundance of different small RNAs, the distribution of piRNAs within the P-element, the length distribution of the piRNAs and the ping-pong signal were computed with previously described Python scripts [51]. For an overview of the composition of all small RNA samples see supplementary table S3.

To identify the positions of piRNA clusters we aligned small RNA reads with a length between 23-29bp to the reference genome of *D. simulans* (strain *w*^*XD*1^ [17]), filtered for uniquely mapping reads (mapping quality >0), counted the number of reads for genomic bins of 500bp, normalized the abundance to a million mapped miRNAs, and clustered bins with a high number of piRNAs using a previously described algorithm that maximizes a local score [26, 51] (*localscore-pirnaclusters.py* --threshold 10 --binsize 500 --max-bin-score 100)

### Gonadal dysgenesis

In addition to the previously published gonadal dysgenesis assays of the cold invasion at generation 57 [51], we performed gonadal dysgenesis assays at generations 62, 71 and 79 for this work. Gonadal dysgenesis assays were performed as described previously [51]. Briefly we kept fly eggs until eclosion at a temperature of 29°C. Eclosed flies were kept for two days at a 23°C on apple juice agar with live yeast, before dissection in PBS. We used the following classification of ovaries: clearly visible ovarioles or eggs (clear), ovarioles barely visible (weak), ovarioles or eggs could not be detected (absent). The percentage of dysgenic ovaries (HD) was computed as 100 ∗ (absent + (weak/2))/(clear + weak + absent)

### TE abundance and diversity

To avoid inconsistencies we analyzed all Pool-Seq data, including the previously published ones, with the same pipeline. We estimated the abundance and diversity of TEs with DeviaTE [76]. Short reads were aligned with bwa sw (v0.7.4 [54]) to the consensus sequences of TEs in *D. melanogaster* [70] and the sequences of three single copy genes (*traffic jam*, *rpl32* and *rhino*). The abundance of TEs is estimated as the coverage of TEs normalized to the coverage of the single copy genes. DeviaTE also provides the position and frequency of internal deletions of the P-element. The number of reads mapping to the P-element (rpm) and the position of P-element insertions in the genome was determined with PoPoolationTE2 (v1.10.04 [50]). To increase the inner distance, which is a major factor determining the power of PoPoolationTE2, we trimmed reads to a length of 75bp (supplementary table S1). Reads were mapped as single-ends with bwa sw (0.7.4 [54]) to a reference genome consisting of the sequence of the P-element (GenBank:X06779.1 [65]) and a long-read based assembly of *D. simulans* (*w*^*XD*1^ [17]). Using PoPoolationTE2 we restored the paired-end information (*se*2*pe*), generated a ppileup file (*ppileup* --map-qual 15), subsampled the ppileup to a uniform coverage of 15 (*subsamplePpileup* --target-coverage 15), identified signatures of TE insertions (*identifySignatures* --mode separate, --min-count 1, --signature-window fix100), estimated the population frequencies of TE signatures (*frequency*), filtered TE signatures (*filterSignatures* --min-count 1, --max-otherte-count 2, --max-structvar-count 2) and finally paired TE signatures to obtain a list of P-element insertions (*pairupSignatures*). To find the positions of ORCs in *D. simulans* we aligned the ORC sequences of *D. melanogaster* [74] to *D. simulans* (*w*^*XD*1^) with bwa sw (v0.7.17; see also [51]). To identify the location of insertions sites shared between *D. melanogaster* and *D. simulans* we extracted 1000bp flanking the previously described insertion sites [50] and aligned these flanking sequences to *D. simulans* (*w*^*XD*1^) with bwa sw (v0.7.17). Finally we filtered for a minimum size of 500 (for ORCs or shared regions) and merged overlapping annotations. The enrichment of P-element insertions (*e*) was computed as *e* = *o*/(*f ∗ t*), where *o* is the observed number of P-element insertion in a feature (ORCs, piRNA clusters, conserved insertion sites), *f* the genomic fraction of the feature and *t* the total number of P-element insertions.

### Telomeric associated sequences (TAS)

To identify the position of TAS we extracted the sequences of all genes of *D. melanogaster* (v6.39; https://flybase.org/) and *D. simulans* (v2.02; https://flybase.org/) into a fasta file and aligned them to the reference genome of *D. simulans* (*w*^*XD*1^) with bwa sw (v0.7.17). We annotated the sequences between the last genes (either from *D. melanogaster* or *D. simulans*) and the telomeric ends of the chromosome arms as TAS (supplementary table S6). To obtain the sequences of anchor reads, i.e. reads where solely the mate but not the focal read maps to the P-element, we aligned all Pool-Seq data to the sequence of the P-element (GenBank: X06779.1) with bwa sw (v0.7.4) and extracted the anchor reads using a custom script (*bam*2*pelematefastq.py*). Finally, the anchor reads were aligned to the TAS sequences, sequences of piRNA clusters or the assembly of *D. simulans* (*w*^*XD*1^) using bwa sw (v0.7.17).

### Simulated TE invasions

Simulations of TE invasions with piRNA clusters were performed with Invade (v0.8.07) [47]. We simulated five chromosome arms of 32.4*Mb* and a piRNA cluster of size 850*kb* at the end of each chromosome giving a total genome size of 162MB and a total size of piRNA clusters of 4.25Mb. We used a population size of *N* = 291 (see [4]) and a uniform recombination rate of *r* = 4*cM/Mb* (with a window size of 2.5Mb the recombination rate in *D. simulans* varies largely between 2.5-5cM/Mb [38]). To trigger the invasions we randomly distributed 498 TE insertions in the population each with a population frequency of 1/2 ∗ 291, which corresponds to 0.86 TE insertions per haploid genome as found in our base populations (supplementary table S2). For hot conditions we used an average transposition rate of *u* = 0.154 and for cold conditions *u* = 0.051 (corresponding to the average transposition rate across the replicates; supplementary table S2). Simulations were run for 1000 generations.

## Author contributions

RK and CS conceived this work. VN prepared sequencing libraries. RK analyzed the data and performed gonadal dysgenesis assays. RK and CS wrote the manuscript.

## Data availability

All scripts used in this work are available at https://sourceforge.net/p/te-tools/code/HEAD/tree/ ( folder “coldinvasion”). All data have been made publicly available at SRA (xxx).

## Acknowledgments

We thank all members of the Institute of Population Genetics for feedback and support. This work was supported by an Austrian Science Foundation (FWF) grant (P29016-B25) to RK and an European Research Council (ERC) Grant “ArchAdapt” to CS.

