## supplementary figures and table for "The transposition rate has little influence on equilibrium copy numbers of the P-element"

### Supplementary figures and tables

September 20, 2021

#### Supplementary Figures

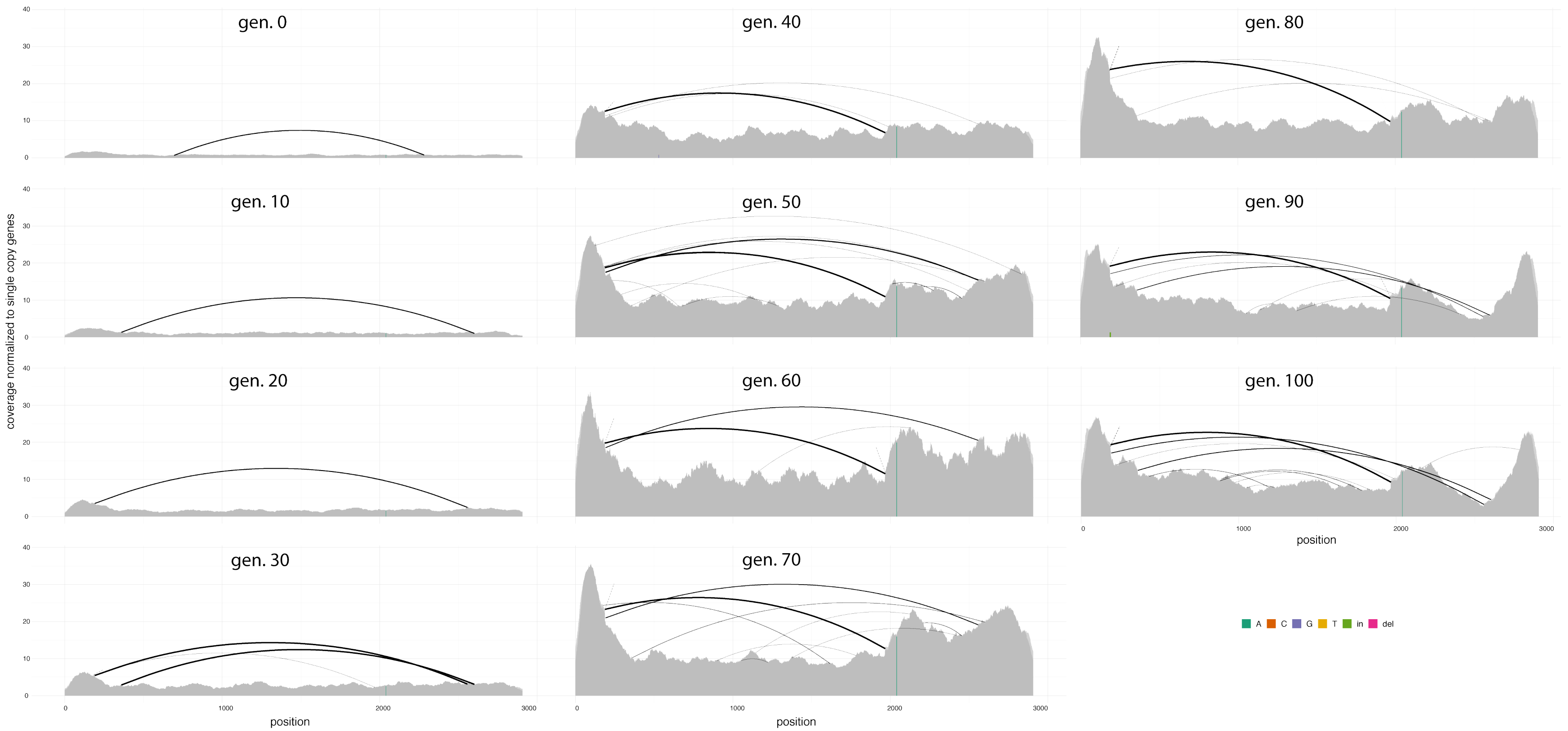


Figure 1: Abundance and diversity of the P-element at cold conditions in replicate 1. Single-nucleotide polymorphisms (SNPs) and small internal deletions (indels) are shown as colored lines. Large internal deletions are shown as black arcs.

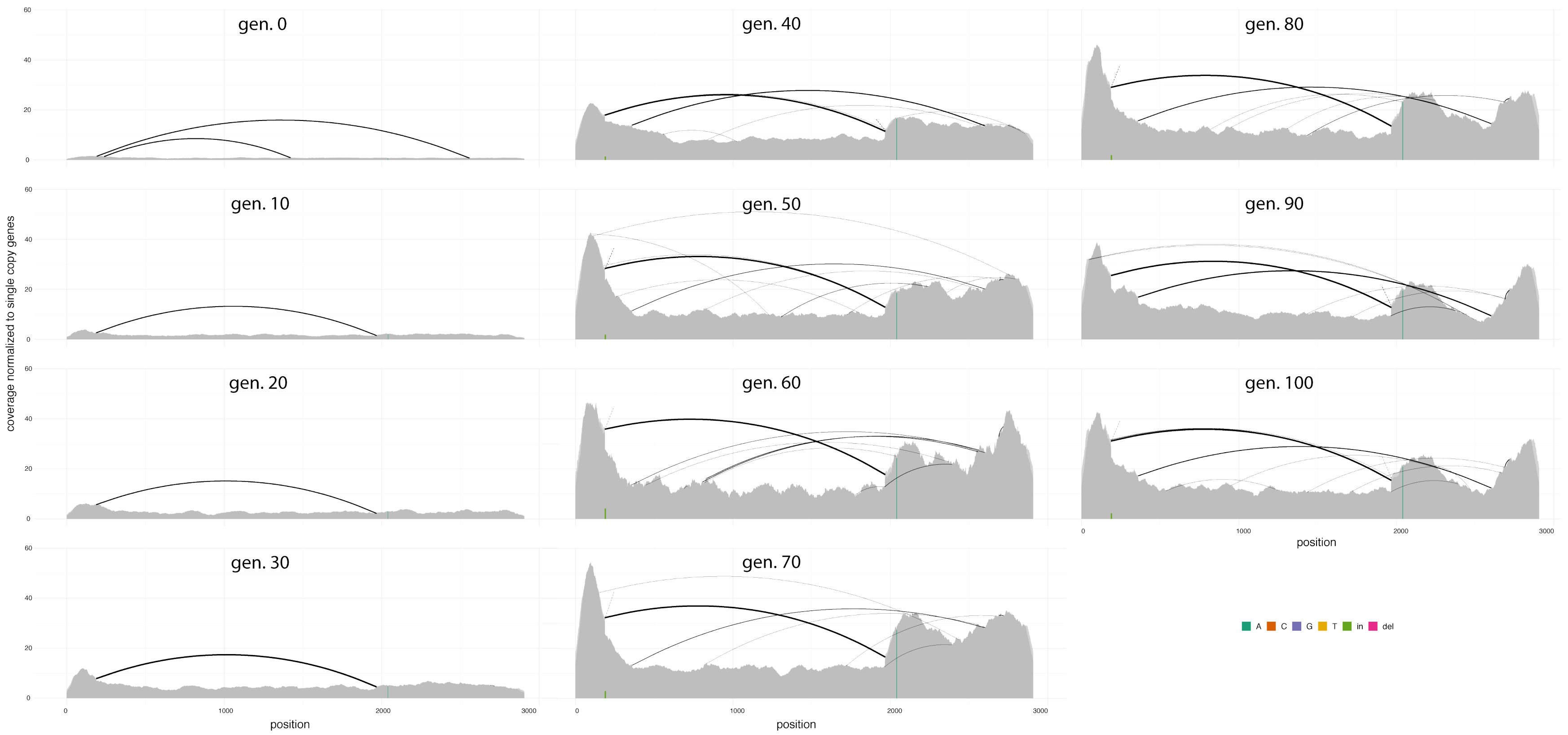


Figure 2: Abundance and diversity of the P-element at cold conditions in replicate 3.

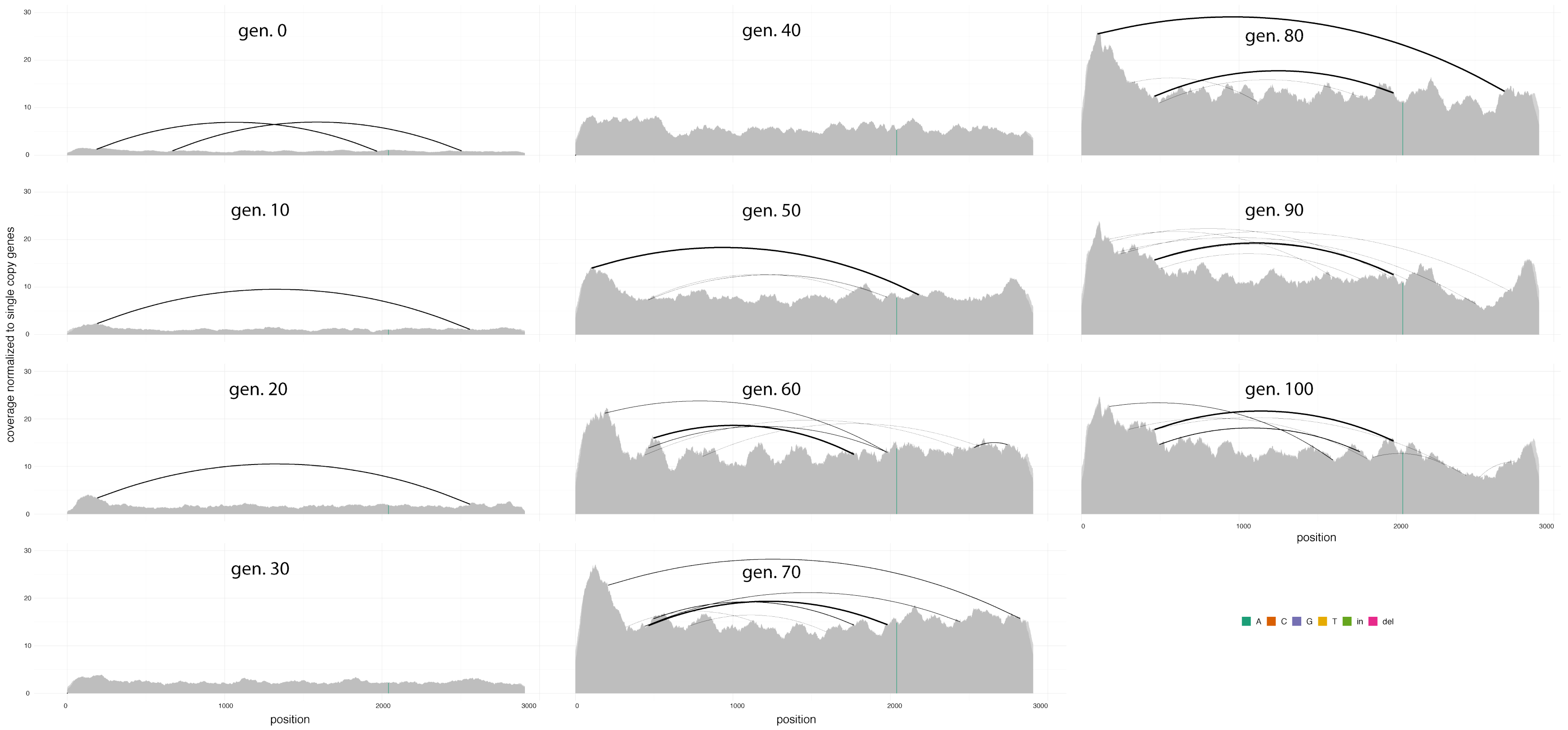


Figure 3: Abundance and diversity of the P-element at cold conditions in replicate 5.

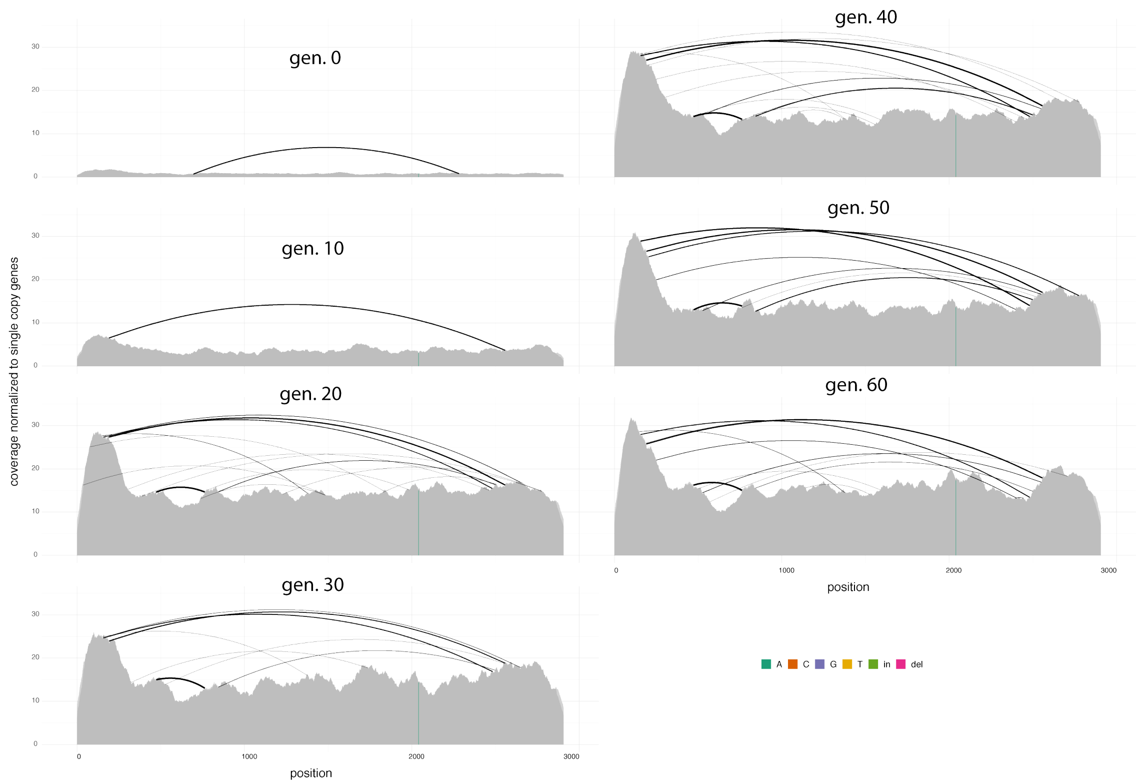


Figure 4: Abundance and diversity of the P-element at hot conditions in replicate 1.

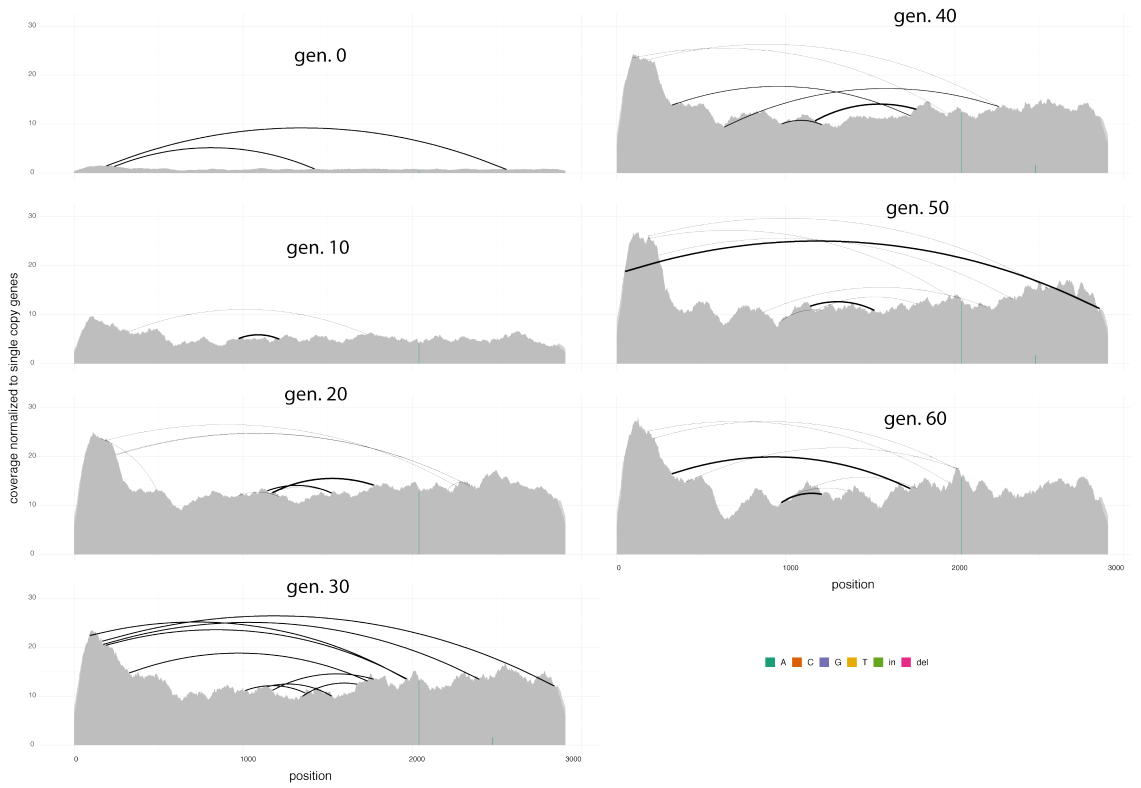


Figure 5: Abundance and diversity of the P-element at hot conditions in replicate 3.

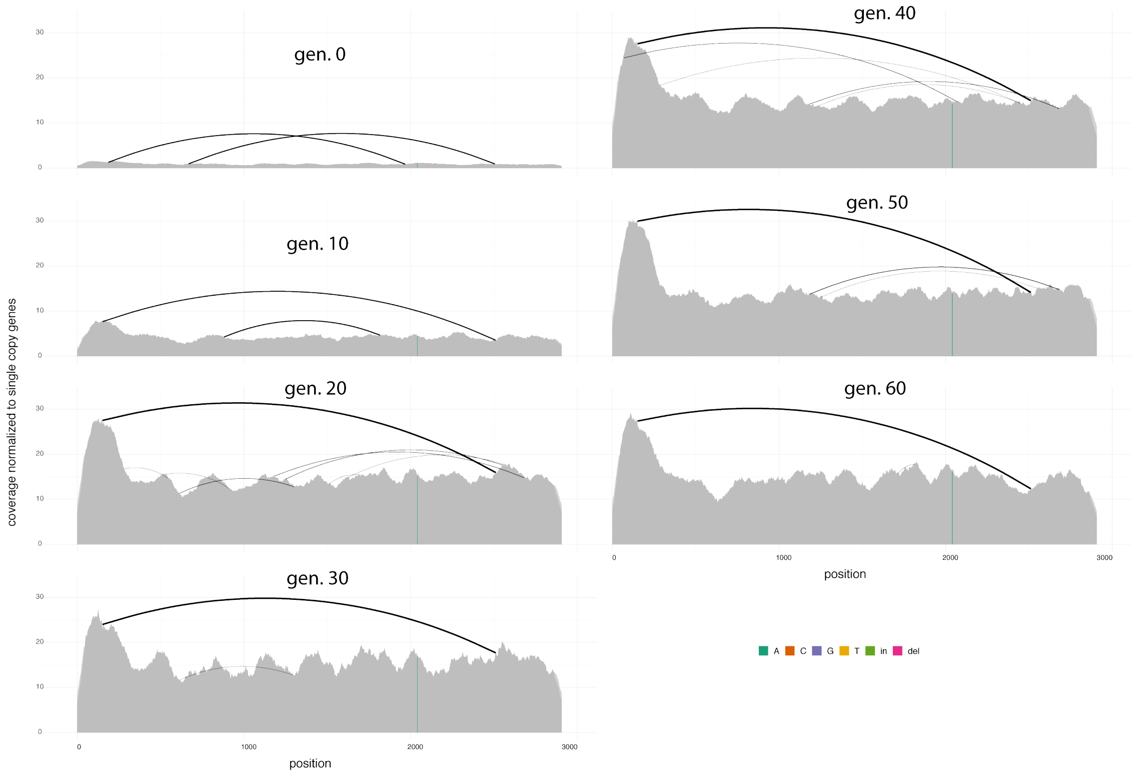


Figure 6: Abundance and diversity of the P-element at hot conditions in replicate 5.

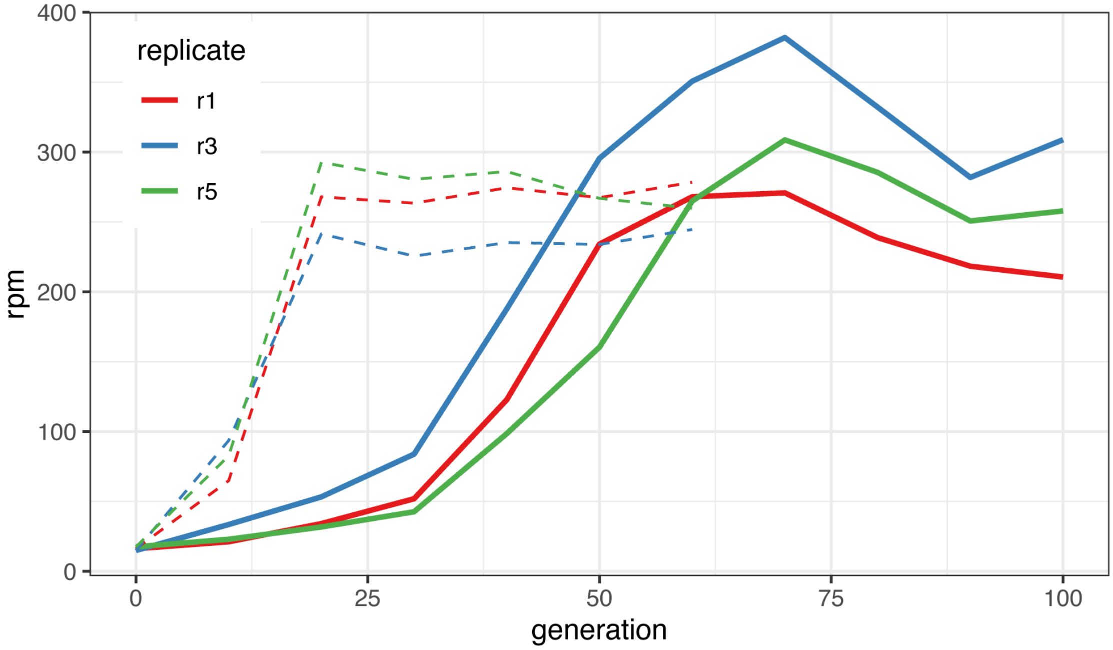


Figure 7: Abundance of P-element insertions during the invasion in three replicates. Here we provide raw estimates of the TE abundance in reads mapping to the P-element out of a million mapped reads (rpm). Hatched line shows the previously published invasion dynamics at hot conditions [Kofler et al., 2018].

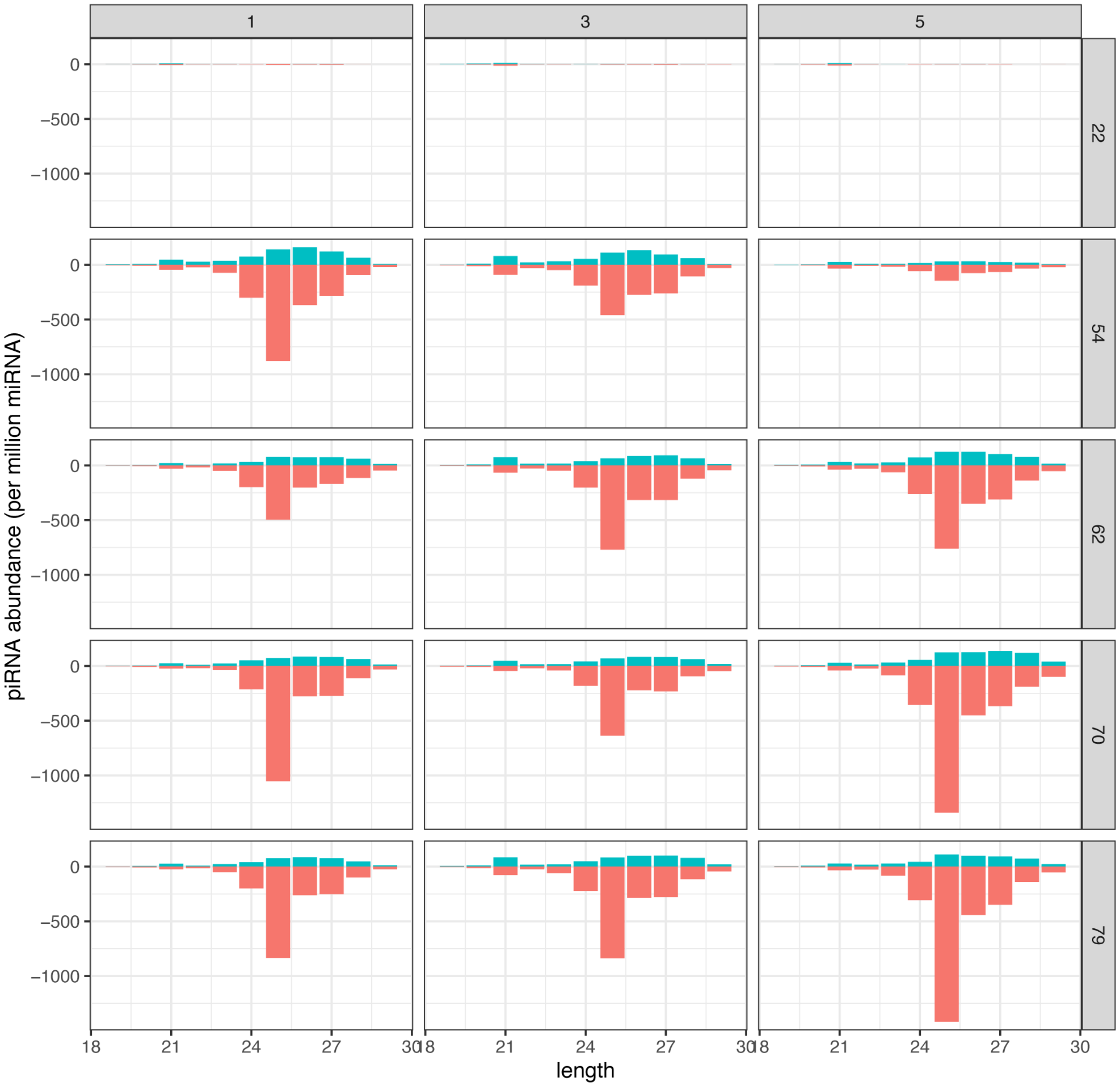


Figure 8: Length distribution of small RNAs (19-29nt) mapping to the P-element. The total abundance of small RNAs was normalized to 1 million miRNAs. Sense RNAs are on the positive y-axis (green) and antisense RNAs on the negative y-axis (red). The replicates are shown in the top panels and the generations in the right panels

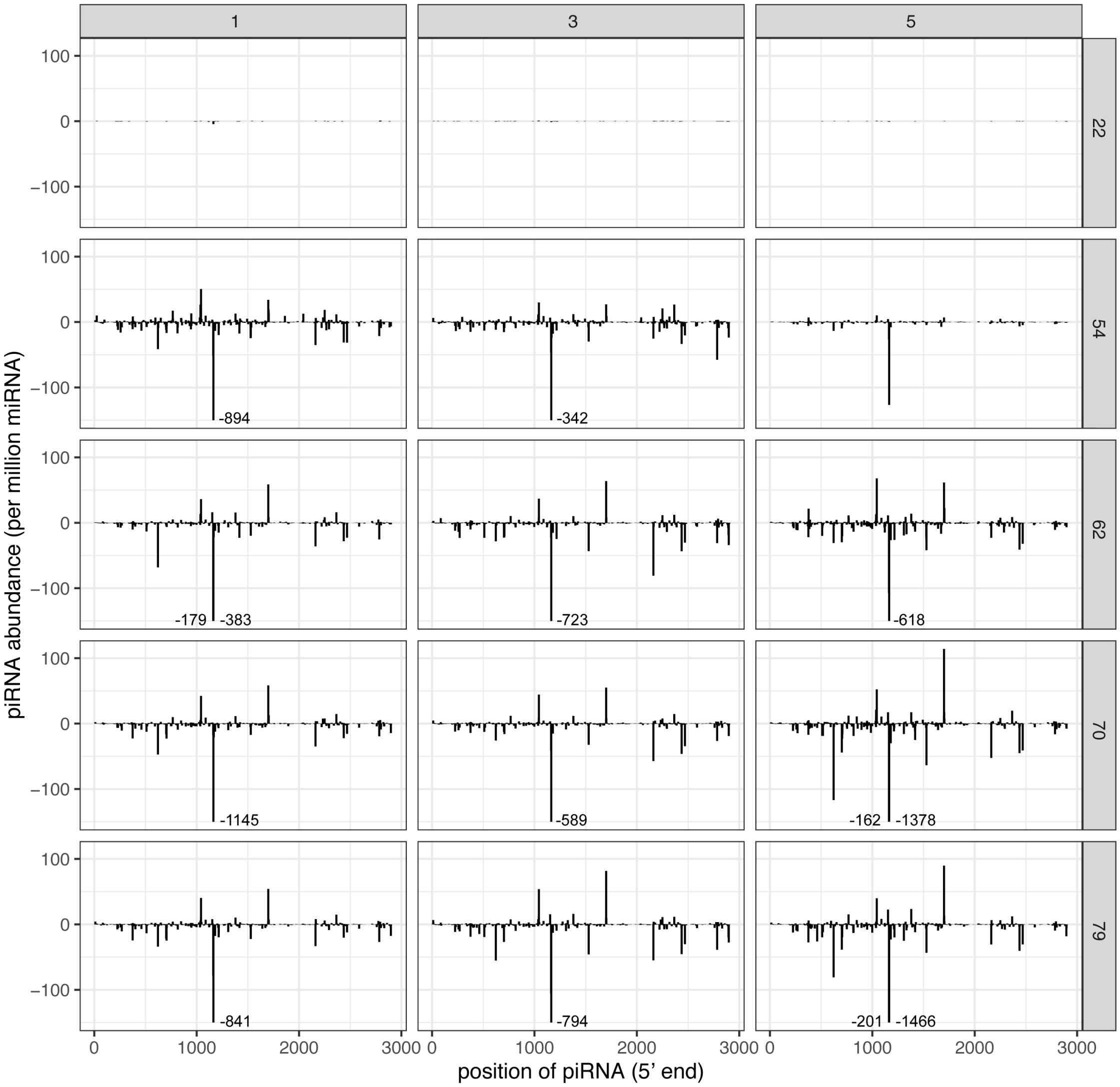


Figure 9: Distribution of piRNAs (23-29nt) along the P-element. Only the 5' positions of piRNAs are considered and piRNA abundance is normalized to one million miRNAs (ppm). Replicates are at the top panel and the generations at the right panel. Sense piRNAs are shown on the positive y-axis and antisense piRNAs on the negative y-axis. Large peaks were truncated (the total abundance of piRNAs is indicated)

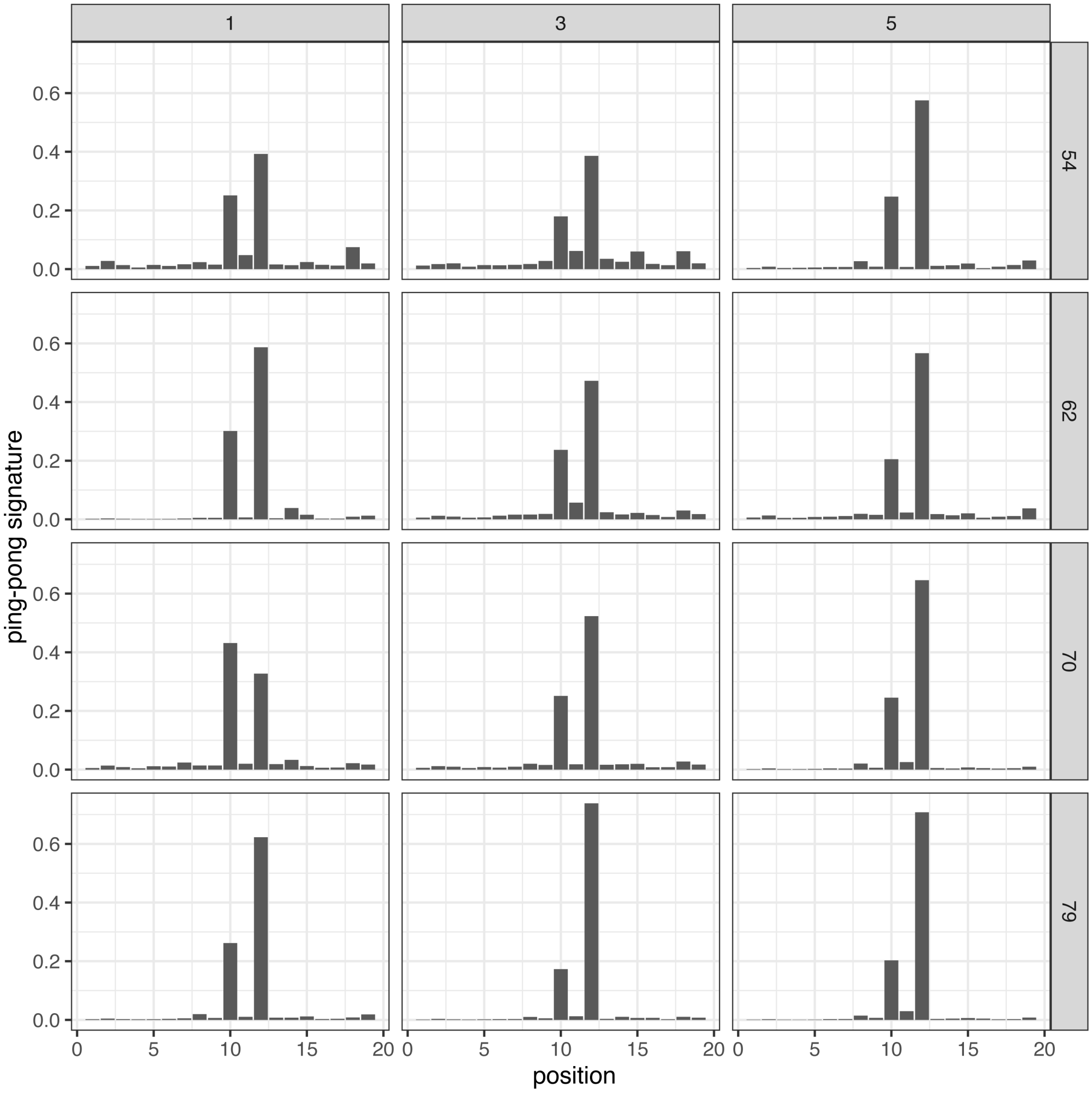


Figure 10: Ping-pong signature of P-element piRNAs. Data are shown for the three replicates (top panel) at different generations during the experiment (right panel). Due to a low number of piRNAs we could not compute ping-pong signatures for any replicate at generation 22.

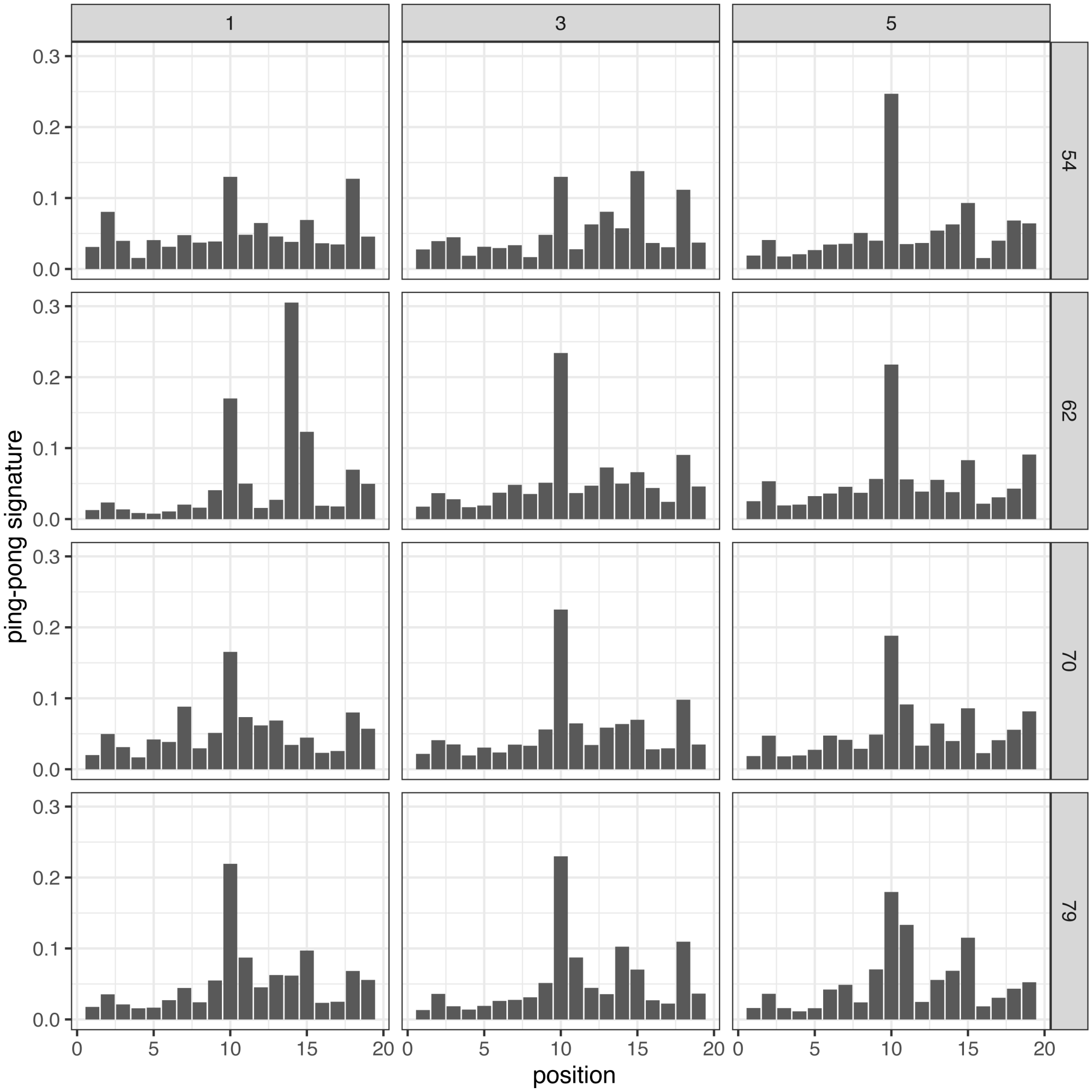


Figure 11: Ping-pong signature of P-element piRNAs during the cold invasion. piRNAs mapping to positions 1162 and 1164 (5'-end) of the P-element are excluded. Data are shown for the three replicates (top panel) at different generations during the experiment (right panel). Due to a low number of piRNAs we could not compute ping-pong signatures for any replicate at generation 22.

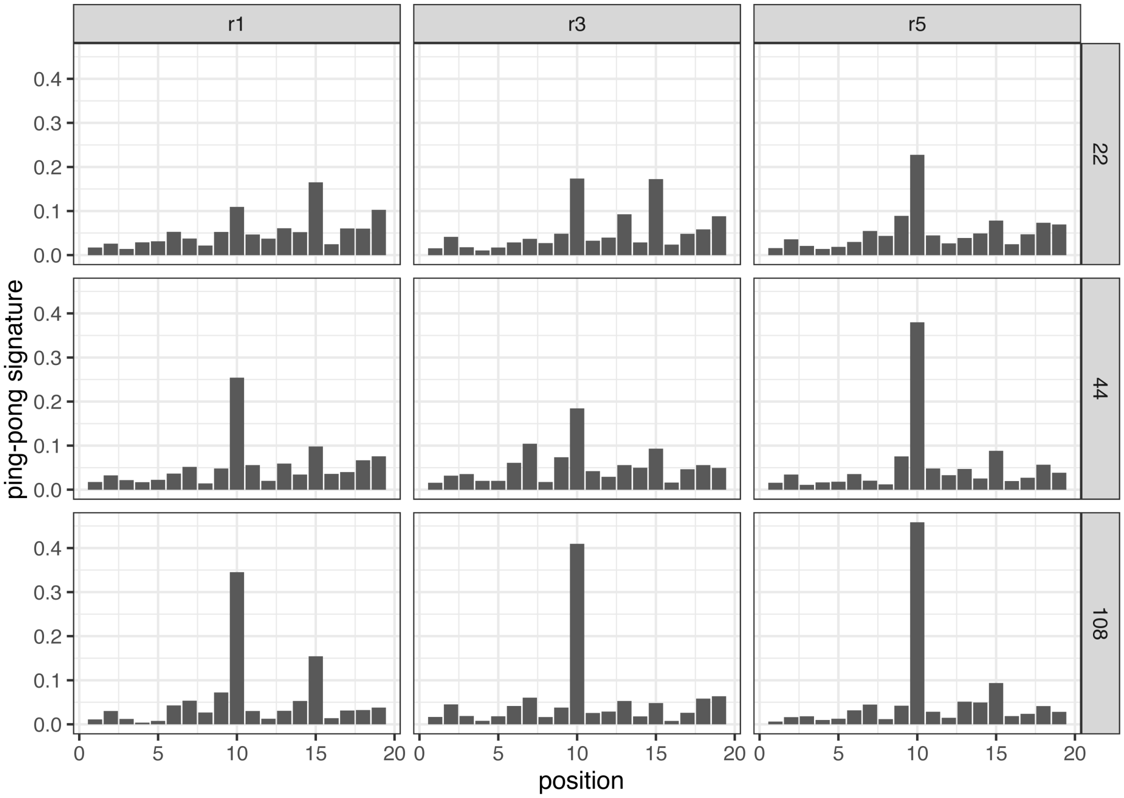


Figure 12: Ping-pong signature of P-element piRNAs during the hot invasion. piRNAs mapping to positions 1162 and 1164 (5'-end) of the P-element are excluded. Data are shown for the three replicates (top panel) at different generations during the experiment (right panel). Note that the ping-pong signature (at position 10) increases during the experiment.

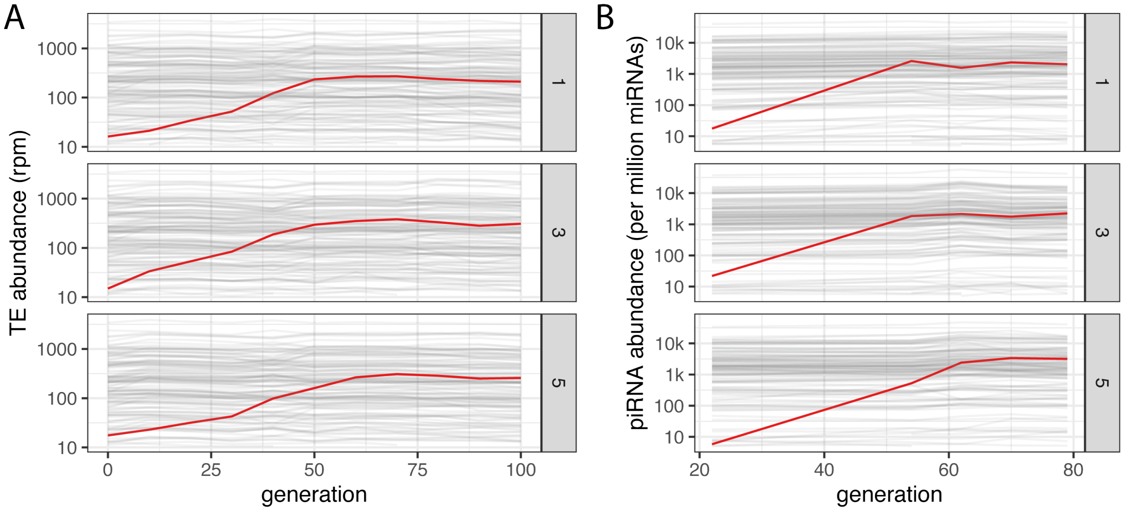


Figure 13: Abundance of TEs and piRNAs during the P-element (red) invasion. Data are shown for 121 TE families (grey; 23 TIR, 60 LTR, 37 non-LTR and 1 Foldback). A) Abundance of TEs in reads mapping to a TE per million reads (rpm). The base population consists solely of females whereas the evolved populations are a mix of males and females. Families with fewer than 10 rpm are not shown. B) Abundance of piRNAs in piRNAs per million miRNAs (ppm). Families with fewer than 5 ppm are not shown. Note that solely the P-element shows a marked increase in copy numbers and piRNAs.

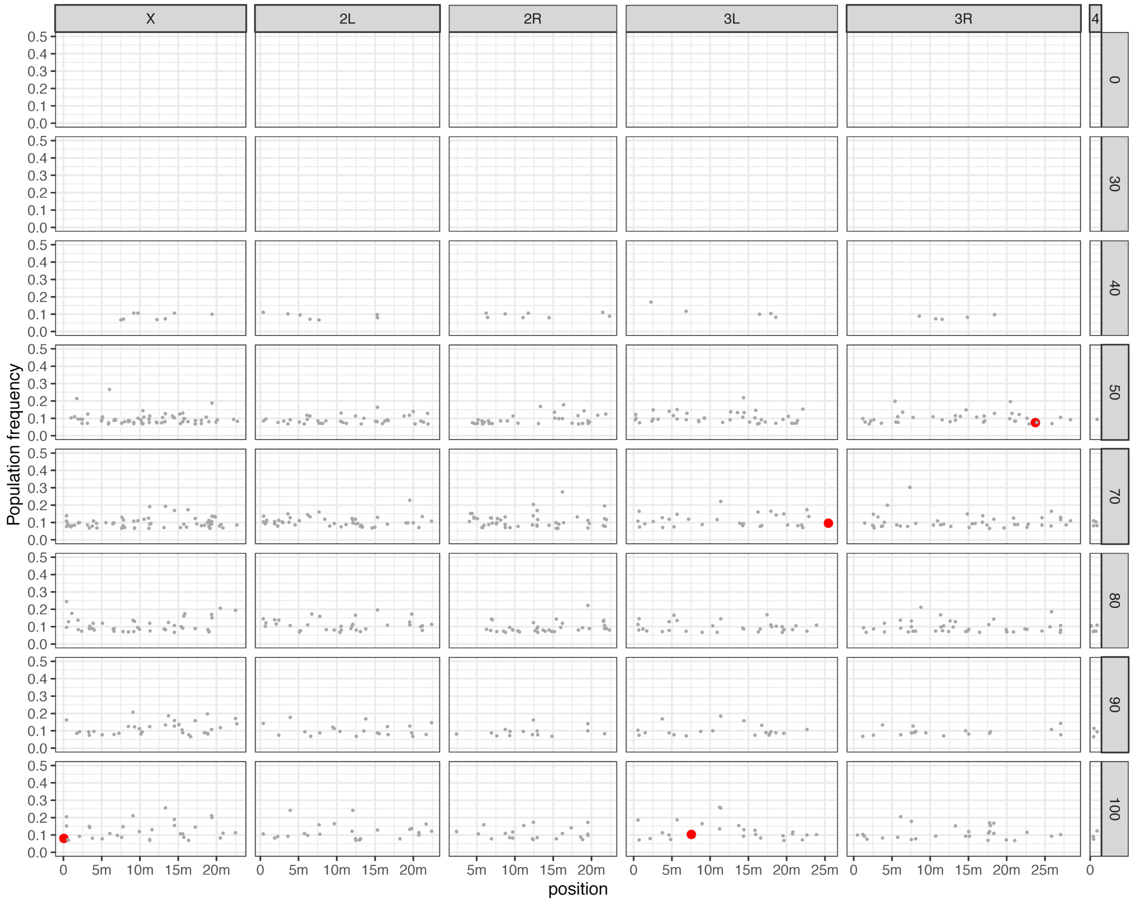


Figure 14: Positions and population frequencies of P-element insertions at cold conditions in replicate 1. The generations are in the right panel and P-element insertions in piRNA-clusters are shown red. Generations 10 and 20 are not shown due to the absence of detectable P-element insertions. To facilitate an unbiased comparison among samples we subsampled the physical coverage to 15. For generation 60 the physical coverage was insufficient. m: million base pairs

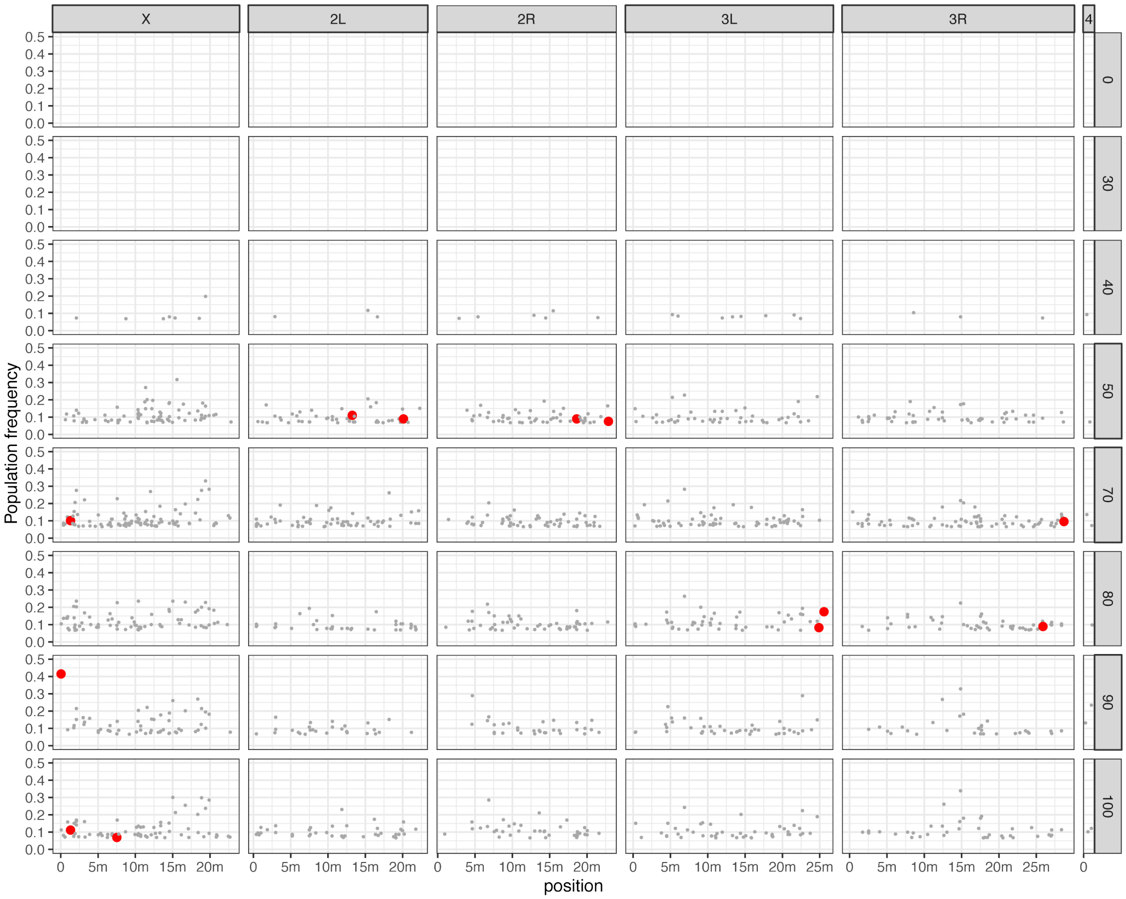


Figure 15: Positions and population frequencies of P-element insertions at cold conditions in replicate 3. For details see supplementary figure 14.

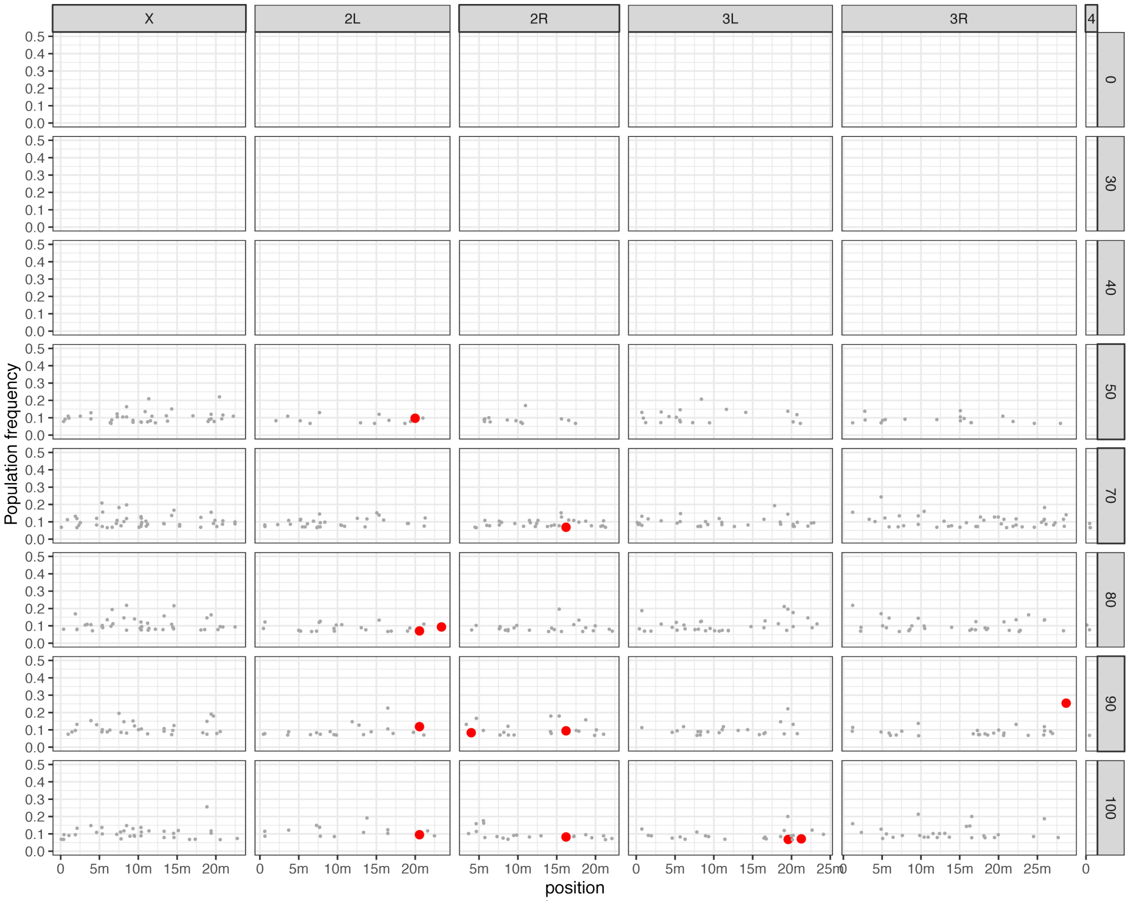


Figure 16: Positions and population frequencies of P-element insertions at cold conditions in replicate 5. For details see supplementary figure 14.

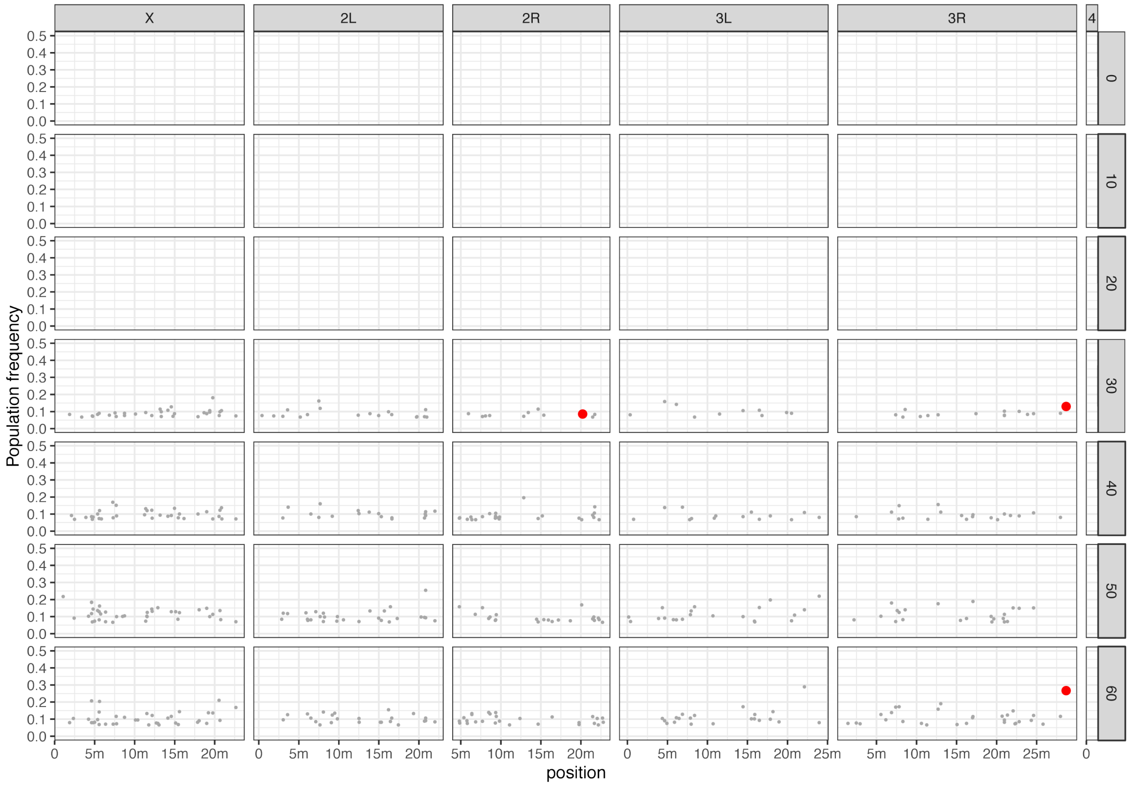


Figure 17: Positions and population frequencies of P-element insertions at hot conditions in replicate 1. The generations are in the right panel and P-element insertions in piRNA-clusters are shown red. We reanalyzed the data of our previous work [Kofler et al., 2018] with the slightly different pipeline used in this work (e.g. different read-length and reference genome) to facilitate a comparison of the P-element invasions at hot and cold conditions. The physical coverage was subsampled to 15. m: million base pairs

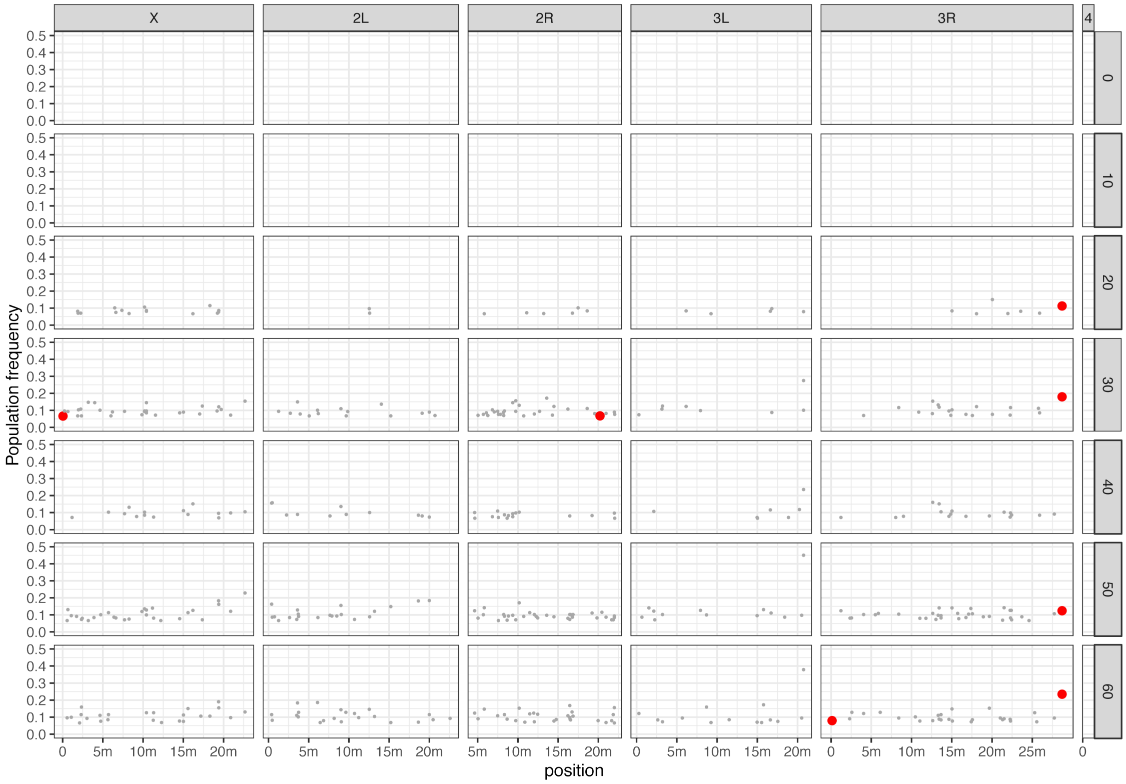


Figure 18: Positions and population frequencies of P-element insertions at hot conditions in replicate 3. For details see supplementary figure 17.

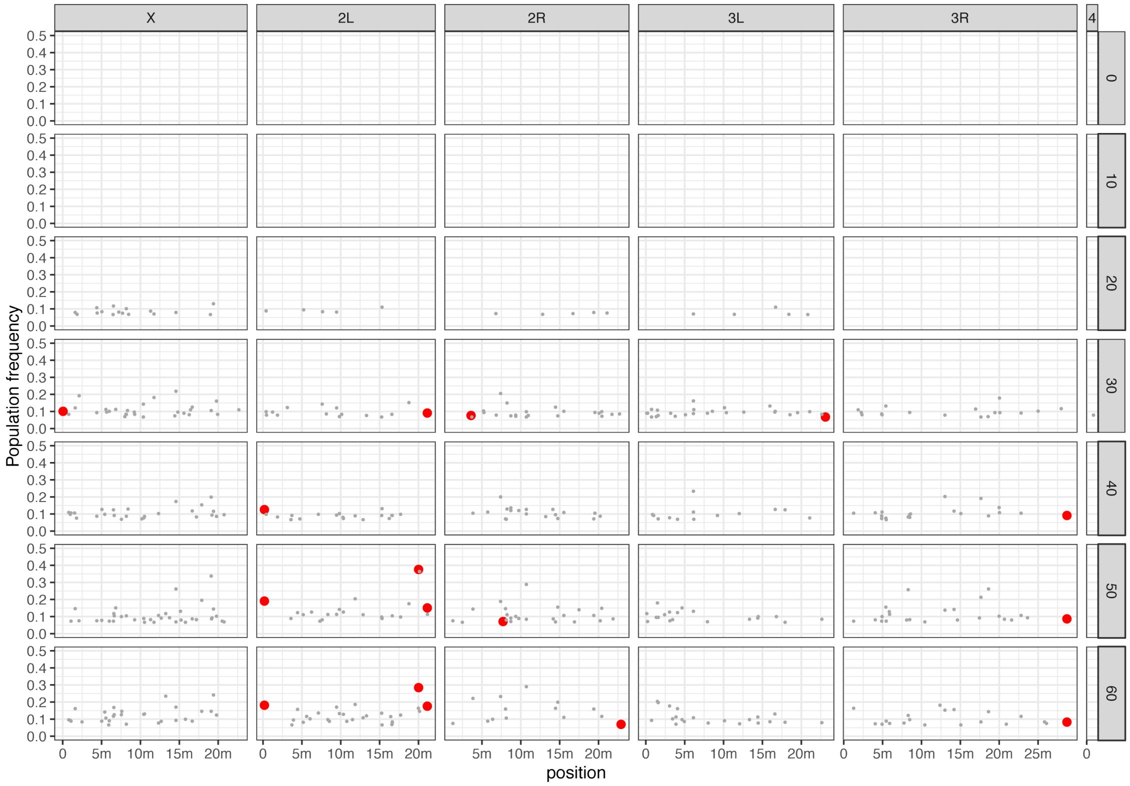


Figure 19: Positions and population frequencies of P-element insertions at hot conditions in replicate 5. For details see supplementary figure 17.

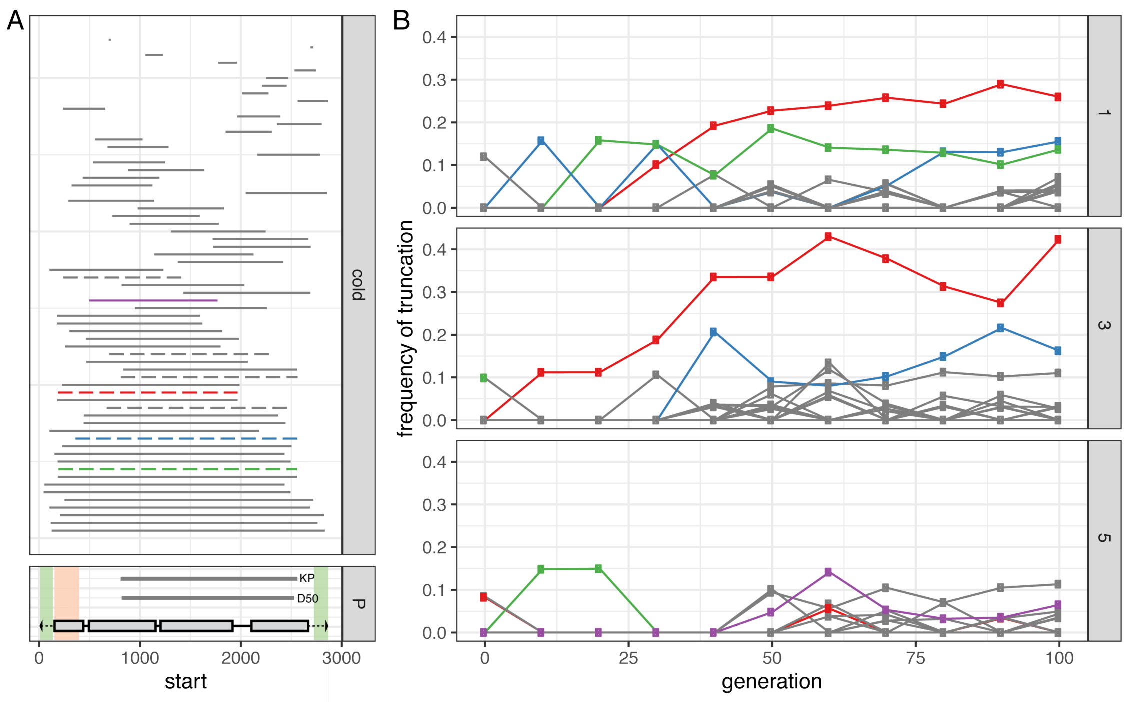


Figure 20: Dynamics of internally deleted P-elements during the invasion A) Positions of all internal deletions (IDs) observed during the cold invasion. Horizontal bars represents the deleted sequence. IDs that were likely present in the base population are shown as dashed line. The lower panel shows the structure of the P-element with the four ORFs (grey box) and the TIRs (black triangles). The position of two IDs that repress P-element activity (KP and D50) are indicated [Black et al., 1987, Rasmusson et al., 1993]. Regions required for mobilization of the P-element are shaded in green, and regions required for repressing P-element activity are shaded in orange [Majumdar and Rio, 2015]. B) Frequency of IDs during the invasion for all three replicates (right panel). The color of the IDs is identical to panel A.

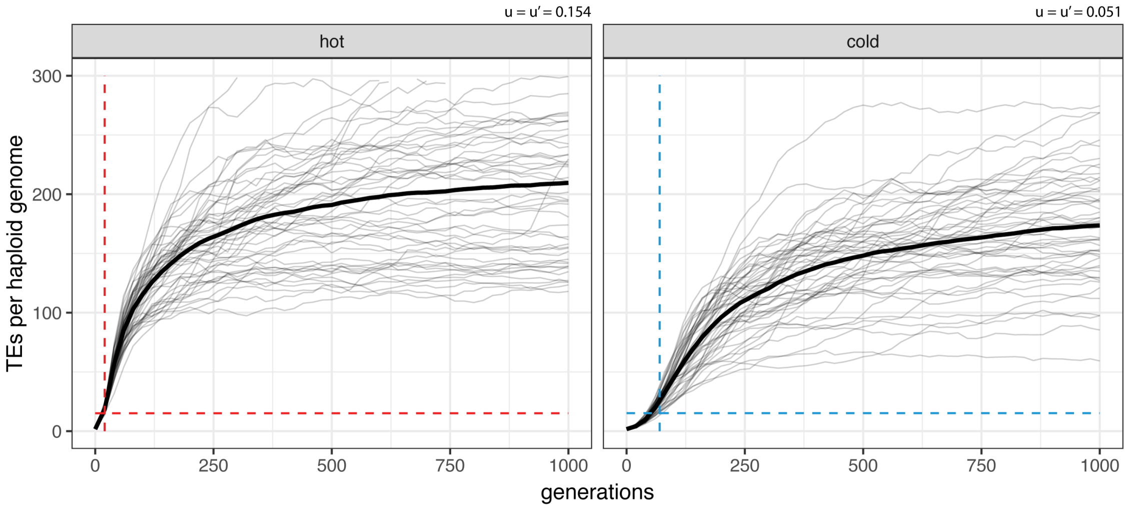


Figure 21: Expected and observed invasion dynamics under the trap model. The solid lines show the expected invasion dynamics with the parameters inferred from the data. Fifty replicates where used and the bold lines are averages. The dashed lines indicate the observed onset and level of the plateau in our experimental populations. Since neutral TE insertions were simulated ($x = 0$), the transposition rate ($u$) is identical to the effective transposition rate ($u' = u - x$).

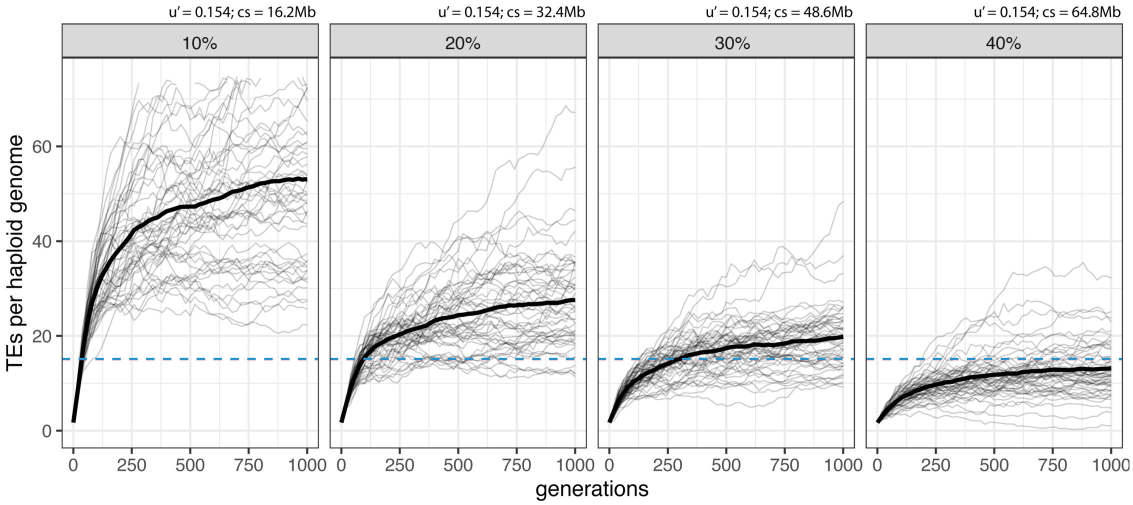


Figure 22: Influence of the size of piRNA clusters on the invasion dynamics. The solid lines show the expected invasion dynamics under the trap model. Fifty replicates where used and the bold lines are the average. The dashed lines indicate the plateau level in our experimental populations. Since neutral TE insertions were simulated, the transposition rate is identical to the effective transposition rate ($u' = u$). $u'$ effective transposition rate, $cs$ cluster size.

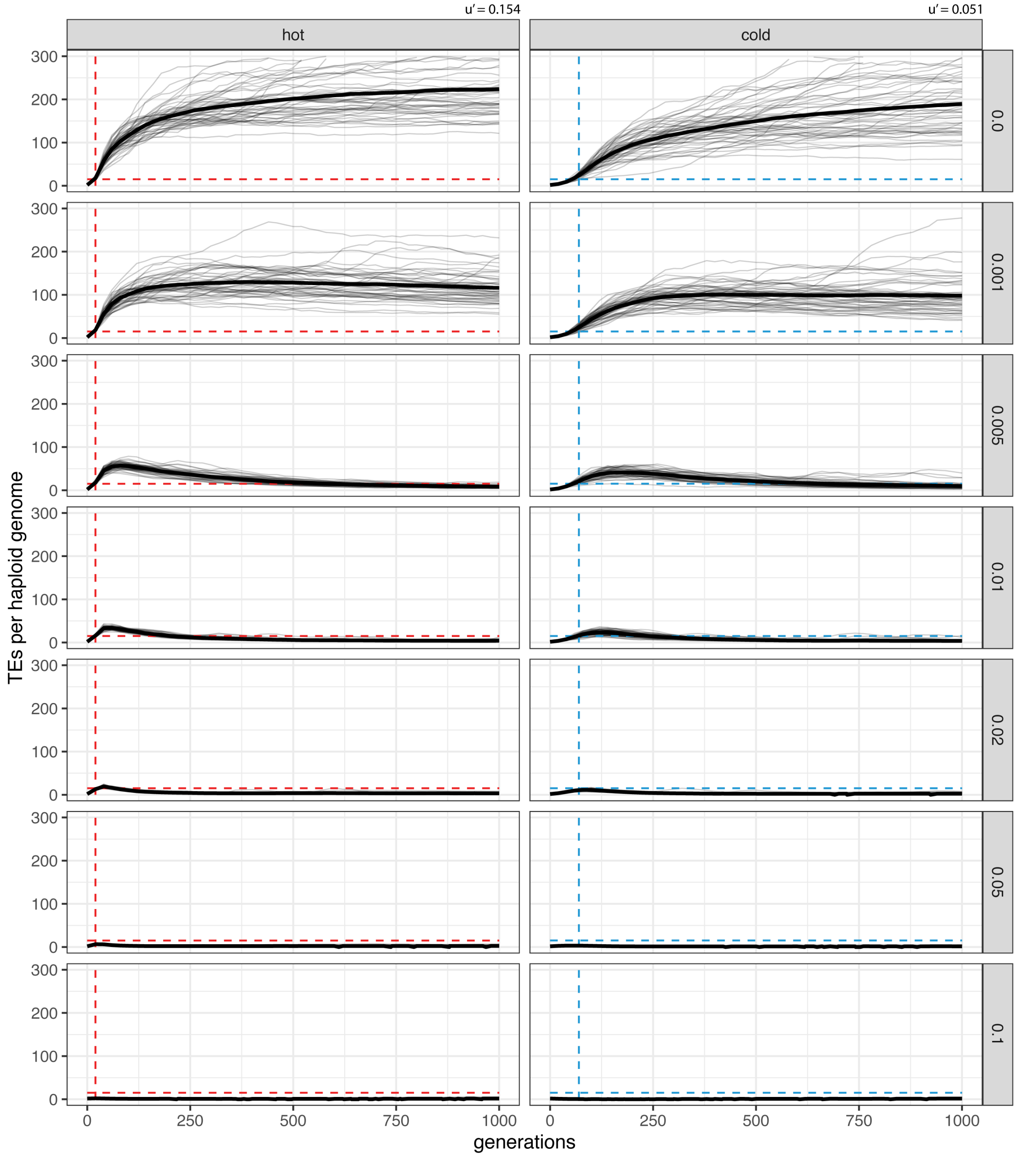


Figure 23: Expected and observed invasion dynamics under the trap model with constant negative effects of TE insertions (insertions in piRNA clusters were neutral). The effective transposition rate ($u' = u - x$) was kept constant by simultaneously changing the negative effect of TE insertions ($x$; right panel) and the transposition rate ($u$) . The solid lines show the expected invasion dynamics. Fifty replicates where used and the bold lines are averages. The dashed lines indicate the observed onset and level of the plateau in our experimental populations. $u'$ effective transposition rate

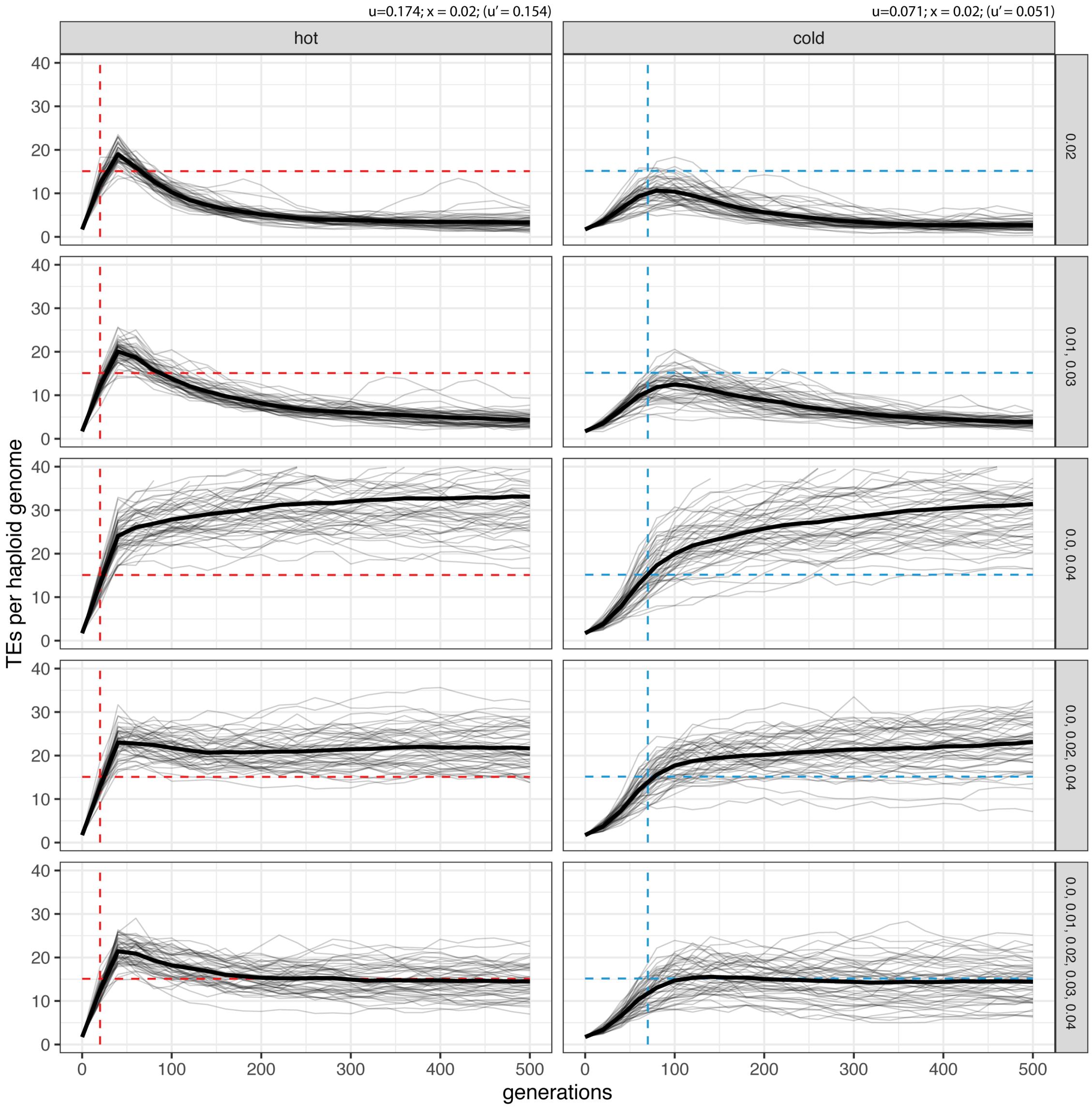


Figure 24: Expected and observed invasion dynamics under the trap model with a distribution of negative effects of TE insertions (the average negative effect was $\bar{x} = 0.02$; insertions in piRNA clusters were neutral). TE insertions had equal proportions of the negative effects indicated in the right panel. For example with "0.0, 0.02, 0.04", about 33.3% of the insertions had a negative effect of $x = 0.0$, another 33.3% had $x = 0.02$, and 33% had $x = 0.04$. The solid lines show the expected invasion dynamics under the trap model. Fifty replicates where used and the bold lines are averages. The dashed lines indicate the observed onset and level of the plateau in our experimental populations. $u'$ effective transposition rate, $cs$ cluster size.

Table 1: Overview of the Pool-Seq data used for estimating the status of the P-element invasion. We show the IDs of the lanes, the read length (rl.), the inner distance (id.) when reads are trimmed to length 75bp as used for finding the insertion sites with PoPoolationTE2 [Kofler et al., 2016], the number of sequenced paired end reads (in millions) for each replicate and the source publication of the data. Reads of several samples (e.g. 313a) were trimmed to a length of 100bp.

| | | | | . | replicate | | | |
|---|---|---|---|---|---|---|---|---|
| sample | lane IDs | Sequencer | rl. | id. | 1 | 3 | 5 | source |
| base | 126a,b,c,d,e | HiSeq 2000 | 100 | 237 | 73.5 | 105.8 | 122.1 | Kofler et al. [2018] |
| hot10 | 127a,b | HiSeq 2000 | 100 | 241 | 34.1 | 27.0 | 44.7 | Kofler et al. [2018] |
| hot20 | 128a,b,c,d | HiSeq 2000 | 100 | 183 | 88.0 | 85.0 | 79.0 | Kofler et al. [2018] |
| hot30 | 129b,c,d | HiSeq 2000 | 100 | 209 | 40.8 | 36.5 | 22.2 | Kofler et al. [2018] |
| hot40 | 130b,c,d | HiSeq 2000 | 100 | 186 | 59.0 | 75.9 | 50.4 | Kofler et al. [2018] |
| hot50 | 131b,c | HiSeq 2000 | 100 | 181 | 29.3 | 30.0 | 36.5 | Kofler et al. [2018] |
| hot60 | 140b,c | HiSeq 2000 | 100 | 236 | 48.6 | 27.9 | 31.6 | Kofler et al. [2018] |
| cold10 | 132b,c | HiSeq 2000 | 100 | 197 | 34.7 | 40.6 | 34.2 | Kofler et al. [2018] |
| cold20 | 133b,c | HiSeq 2000 | 100 | 196 | 42.7 | 25.0 | 24.4 | Kofler et al. [2018] |
| cold30 | 134b,c | HiSeq 2000 | 100 | 183 | 29.2 | 32.7 | 33.1 | Kofler et al. [2018] |
| cold40 | 185a,b,c | HiSeq 2500 | 120(100) | 360 | 19.4 | 46.9 | 21.7 | Kofler et al. [2018] |
| cold50 | 313a | HiSeq X Ten | 150(100) | 131 | 38.7 | 41.0 | 38.8 | this work |
| cold60 | 329a | HiSeq X Ten | 150(100) | 114 | 13.3 | 12.2 | 33.0 | this work |
| cold70 | 393a | HiSeq X Ten | 150(100) | 108 | 39.9 | 45.3 | 50.9 | this work |
| cold80 | 487a | HiSeq X Ten | 150(100) | 128 | 34.7 | 37.4 | 42.5 | this work |
| cold90 | 542a | HiSeq X Ten | 150(100) | 173 | 40.0 | 42.0 | 40.0 | this work |
| cold100 | 543a | HiSeq X Ten | 150(100) | 174 | 41.0 | 39.8 | 37.9 | this work |

### Supplementary Tables

Table 2: Abundance and effective transposition rate ($u'$) of the P-element in experimental evolving populations. Data are shown for different temperatures (temp), generations (gen) and replicates (columns 1, 3, 5). The abundance of the P-elements was estimated as reads per million (rpm) using PoPoolationTE2 [Kofler et al., 2016] and as insertions per haploid genome (ins) using DeviaTE [Weilguny and Kofler, 2019].

| | | rpm | | | ins | | | $u'_{rpm}$ | | | $u'_{ins}$ | | |
|---|---|---|---|---|---|---|---|---|---|---|---|---|---|
| temp | gen | 1 | 3 | 5 | 1 | 3 | 5 | 1 | 3 | 5 | 1 | 3 | 5 |
| base | 0 | 16.2 | 14.9 | 17.5 | 0.85 | 0.80 | 0.92 | - | - | - | - | - | - |
| hot | 10 | 64.9 | 93.6 | 82.6 | 3.89 | 5.32 | 4.38 | 0.149 | 0.201 | 0.168 | 0.164 | 0.209 | 0.169 |
| hot | 20 | 268.0 | 241.5 | 292.7 | 15.73 | 13.68 | 15.59 | 0.152 | 0.099 | 0.135 | 0.150 | 0.099 | 0.135 |
| hot | 30 | 263.4 | 225.5 | 280.5 | 15.94 | 13.08 | 16.20 | -0.002 | -0.007 | -0.004 | 0.001 | -0.004 | 0.004 |
| hot | 40 | 274.4 | 235.2 | 286.1 | 15.69 | 13.09 | 16.24 | 0.004 | 0.004 | 0.002 | -0.002 | 0.000 | 0.000 |
| hot | 50 | 267.4 | 233.9 | 266.9 | 15.74 | 13.32 | 15.04 | -0.003 | -0.001 | -0.007 | 0.000 | 0.002 | -0.008 |
| hot | 60 | 278.4 | 244.6 | 259.8 | 17.35 | 14.04 | 15.64 | 0.004 | 0.004 | -0.003 | 0.010 | 0.005 | 0.004 |
| cold | 10 | 21.2 | 33.5 | 22.9 | 1.21 | 1.83 | 1.20 | 0.027 | 0.084 | 0.027 | 0.036 | 0.086 | 0.027 |
| cold | 20 | 34.1 | 53.3 | 31.8 | 1.87 | 2.88 | 1.81 | 0.049 | 0.048 | 0.033 | 0.045 | 0.047 | 0.042 |
| cold | 30 | 51.9 | 83.9 | 42.6 | 3.17 | 5.11 | 2.41 | 0.043 | 0.046 | 0.030 | 0.054 | 0.059 | 0.029 |
| cold | 40 | 122.8 | 187.7 | 98.5 | 7.89 | 12.80 | 5.77 | 0.090 | 0.084 | 0.087 | 0.096 | 0.096 | 0.091 |
| cold | 50 | 234.1 | 295.5 | 160.4 | 12.80 | 16.78 | 8.54 | 0.067 | 0.046 | 0.050 | 0.050 | 0.027 | 0.040 |
| cold | 60 | 268.0 | 350.7 | 264.9 | 15.16 | 22.46 | 13.55 | 0.014 | 0.017 | 0.051 | 0.017 | 0.030 | 0.047 |
| cold | 70 | 270.8 | 381.9 | 308.7 | 15.25 | 21.65 | 15.49 | 0.001 | 0.009 | 0.015 | 0.001 | -0.004 | 0.014 |
| cold | 80 | 238.8 | 332.1 | 285.4 | 12.77 | 18.08 | 13.63 | -0.012 | -0.014 | -0.008 | -0.018 | -0.018 | -0.013 |
| cold | 90 | 218.3 | 281.8 | 250.7 | 11.93 | 16.24 | 12.62 | -0.009 | -0.016 | -0.013 | -0.007 | -0.011 | -0.008 |
| cold | 100 | 210.6 | 308.8 | 257.9 | 12.14 | 18.16 | 13.87 | -0.004 | 0.009 | 0.003 | 0.002 | 0.011 | 0.009 |

Table 3: Overview of small RNA data used in this work. We sequenced small RNA from whole flies at multiple time points during the experimental invasions and assessed the abundance of reads mapping to TEs (i.e. siRNAs and piRNAs), miRNAs, tRNAs, rRNAs and mRNAs. For each class of small RNAs we also estimated the fraction of reads mapping to the sense strand. Samples marked with a star (*) were published in a previous work [Kofler et al., 2018]. temp.: temperature, r.: replicate, gen.: generation, t.[M]: total number of reads in million, m.[M]: mapped reads in million

| | | | | | abundance [%] | | | | | fraction sense[%] | | | | |
|---|---|---|---|---|---|---|---|---|---|---|---|---|---|---|
| temp. | r. | gen. | t.[M] | m.[M] | TE | miRNA | tRNA | rRNA | mRNA | TE | miRNA | tRNA | rRNA | mRNA |
| hot* | 1 | 22 | 14.0 | 7.5 | 22.4 | 58.8 | 8.4 | 1.4 | 9.0 | 25.6 | 100.0 | 100.0 | 99.8 | 61.0 |
| hot* | 3 | 22 | 13.7 | 6.1 | 18.4 | 63.1 | 7.2 | 2.7 | 8.5 | 26.8 | 100.0 | 100.0 | 99.9 | 67.6 |
| hot* | 5 | 22 | 14.6 | 7.6 | 22.2 | 65.7 | 2.7 | 2.3 | 7.1 | 26.4 | 100.0 | 100.0 | 99.8 | 51.4 |
| hot* | 1 | 44 | 13.8 | 6.5 | 15.8 | 40.4 | 21.5 | 9.5 | 12.9 | 23.9 | 100.0 | 100.0 | 100.0 | 81.5 |
| hot* | 3 | 44 | 14.2 | 7.7 | 15.8 | 40.0 | 24.5 | 7.8 | 11.9 | 25.5 | 100.0 | 100.0 | 100.0 | 79.2 |
| hot* | 5 | 44 | 13.6 | 7.1 | 24.6 | 49.7 | 6.9 | 10.6 | 8.2 | 23.6 | 100.0 | 100.0 | 100.0 | 58.4 |
| hot* | 1 | 108 | 12.8 | 6.8 | 27.2 | 60.5 | 1.1 | 4.4 | 6.9 | 25.2 | 100.0 | 99.9 | 99.9 | 35.0 |
| hot* | 3 | 108 | 14.7 | 7.2 | 18.0 | 70.2 | 1.6 | 5.7 | 4.6 | 25.1 | 100.0 | 99.9 | 100.0 | 42.1 |
| hot* | 5 | 108 | 15.2 | 7.6 | 25.9 | 60.0 | 1.1 | 6.5 | 6.5 | 25.0 | 100.0 | 99.8 | 100.0 | 37.3 |
| cold* | 1 | 22 | 12.7 | 5.0 | 17.4 | 51.0 | 11.8 | 9.0 | 10.8 | 24.9 | 100.0 | 100.0 | 100.0 | 76.0 |
| cold* | 3 | 22 | 18.5 | 7.5 | 16.1 | 50.0 | 8.4 | 10.6 | 14.9 | 23.5 | 100.0 | 100.0 | 100.0 | 83.3 |
| cold* | 5 | 22 | 17.7 | 7.3 | 18.7 | 56.3 | 5.8 | 12.0 | 7.2 | 24.5 | 100.0 | 100.0 | 100.0 | 64.3 |
| cold* | 1 | 54 | 12.3 | 6.1 | 25.5 | 61.7 | 1.2 | 5.8 | 5.9 | 24.6 | 100.0 | 99.9 | 99.9 | 40.5 |
| cold* | 3 | 54 | 14.4 | 7.8 | 23.7 | 61.9 | 1.8 | 6.6 | 6.0 | 24.8 | 100.0 | 99.9 | 100.0 | 42.6 |
| cold* | 5 | 54 | 16.7 | 8.5 | 21.8 | 62.8 | 1.8 | 8.2 | 5.3 | 23.1 | 100.0 | 99.9 | 100.0 | 38.2 |
| cold | 1 | 62 | 37.5 | 19.0 | 26.1 | 63.7 | 1.4 | 2.7 | 6.1 | 24.5 | 100.0 | 99.9 | 99.9 | 43.1 |
| cold | 3 | 62 | 46.4 | 24.1 | 29.0 | 59.8 | 0.9 | 3.4 | 7.0 | 24.7 | 100.0 | 99.8 | 99.9 | 37.1 |
| cold | 5 | 62 | 40.9 | 21.2 | 26.7 | 61.0 | 1.1 | 5.1 | 6.1 | 24.4 | 100.0 | 99.8 | 99.9 | 40.0 |
| cold | 1 | 70 | 35.0 | 18.2 | 26.5 | 61.7 | 1.7 | 4.3 | 5.9 | 24.6 | 100.0 | 99.9 | 99.9 | 43.5 |
| cold | 3 | 70 | 44.0 | 23.1 | 24.7 | 63.8 | 1.8 | 3.9 | 5.9 | 25.6 | 100.0 | 99.9 | 99.9 | 41.1 |
| cold | 5 | 70 | 43.1 | 22.1 | 26.2 | 63.2 | 1.5 | 3.0 | 6.1 | 25.0 | 100.0 | 99.9 | 99.9 | 40.0 |
| cold | 1 | 79 | 42.4 | 22.5 | 25.6 | 61.4 | 1.0 | 6.3 | 5.7 | 25.1 | 100.0 | 99.8 | 99.9 | 43.4 |
| cold | 3 | 79 | 39.8 | 19.9 | 24.0 | 64.6 | 1.2 | 4.3 | 5.8 | 25.8 | 100.0 | 99.9 | 99.9 | 40.7 |
| cold | 5 | 79 | 34.3 | 17.2 | 24.2 | 63.6 | 1.2 | 5.3 | 5.7 | 26.2 | 100.0 | 99.9 | 99.9 | 40.9 |

Table 4: Extent of gonadal dysgenesis during the P-element invasion. Results are shown at different generations for the three replicates. We estimated the number of flies having clearly visible ovarioles (normal), weakly visible ovarioles (weak), and no discernible ovarioles (dysgenic) at 29°C. Ovary dissections for each generation were done on a single day. The percentage of dysgenic ovaries (HD) is computed as $100 * (dysgenic + (weak/2))/(normal + weak + dysgenic)$

| generation | replicate | normal | weak | dysgenic | dysgenic [%] |
|---|---|---|---|---|---|
| 57 | 1 | 309 | 18 | 34 | 11.91 |
| 57 | 3 | 241 | 21 | 40 | 16.72 |
| 57 | 5 | 123 | 25 | 154 | 55.13 |
| 62 | 1 | 127 | 1 | 15 | 10.84 |
| 62 | 3 | 154 | 2 | 15 | 9.36 |
| 62 | 5 | 105 | 4 | 64 | 38.15 |
| 71 | 1 | 152 | 4 | 3 | 3.14 |
| 71 | 3 | 135 | 3 | 4 | 3.87 |
| 71 | 5 | 140 | 3 | 7 | 5.67 |
| 79 | 1 | 81 | 0 | 1 | 1.22 |
| 79 | 3 | 56 | 0 | 0 | 0.00 |
| 79 | 5 | 52 | 0 | 1 | 1.89 |

Table 5: Number of TE insertions in piRNA clusters and TAS regions; For each condition (cond), generation (g), replicate (r) we show an unbiased estimate of the number P-element insertions (Pi), the number of cluster insertions with and without TAS insertions (wTAS, woTAS) estimated by PoPoolationTE2, the number of TAS insertions based on anchor reads (anchor TAS; see supplementary table 7) and the sum of cluster and TAS insertions (cti $= woTAS + anchor\_TAS$).

| | | | | PopoolationTE2 | | anchor | |
|---|---|---|---|---|---|---|---|
| cond | g | r | Pi | wTAS | woTAS | TAS | cti |
| base | 0 | 1 | 0.9 | 0.00 | 0.00 | 0.02 | 0.02 |
| base | 0 | 3 | 0.8 | 0.00 | 0.00 | 0.02 | 0.02 |
| base | 0 | 5 | 0.9 | 0.00 | 0.00 | 0.02 | 0.02 |
| cold | 10 | 1 | 1.2 | 0.00 | 0.00 | 0.05 | 0.05 |
| cold | 10 | 3 | 1.8 | 0.00 | 0.00 | 0.03 | 0.03 |
| cold | 10 | 5 | 1.2 | 0.00 | 0.00 | 0.08 | 0.08 |
| cold | 20 | 1 | 1.9 | 0.00 | 0.00 | 0.03 | 0.03 |
| cold | 20 | 3 | 2.9 | 0.00 | 0.00 | 0.05 | 0.05 |
| cold | 20 | 5 | 1.8 | 0.00 | 0.00 | 0.01 | 0.01 |
| cold | 30 | 1 | 3.2 | 0.00 | 0.00 | 0.03 | 0.03 |
| cold | 30 | 3 | 5.1 | 0.00 | 0.00 | 0.11 | 0.11 |
| cold | 30 | 5 | 2.4 | 0.00 | 0.00 | 0.13 | 0.13 |
| cold | 40 | 1 | 7.9 | 0.00 | 0.00 | 0.09 | 0.09 |
| cold | 40 | 3 | 12.8 | 0.00 | 0.00 | 0.24 | 0.24 |
| cold | 40 | 5 | 5.8 | 0.00 | 0.00 | 0.07 | 0.07 |
| cold | 50 | 1 | 12.8 | 0.08 | 0.08 | 0.16 | 0.23 |
| cold | 50 | 3 | 16.8 | 0.36 | 0.36 | 0.22 | 0.58 |
| cold | 50 | 5 | 8.5 | 0.10 | 0.10 | 0.13 | 0.22 |
| cold | 60 | 1 | 15.2 | 0.00 | 0.00 | 0.45 | 0.45 |
| cold | 60 | 3 | 22.5 | 0.00 | 0.00 | 0.27 | 0.27 |
| cold | 60 | 5 | 13.5 | 0.18 | 0.18 | 0.11 | 0.28 |
| cold | 70 | 1 | 15.2 | 0.10 | 0.10 | 0.32 | 0.42 |
| cold | 70 | 3 | 21.6 | 0.20 | 0.10 | 0.36 | 0.46 |
| cold | 70 | 5 | 15.5 | 0.07 | 0.07 | 0.23 | 0.30 |
| cold | 80 | 1 | 12.8 | 0.00 | 0.00 | 0.22 | 0.22 |
| cold | 80 | 3 | 18.1 | 0.35 | 0.35 | 0.22 | 0.57 |
| cold | 80 | 5 | 13.6 | 0.16 | 0.16 | 0.27 | 0.43 |
| cold | 90 | 1 | 11.9 | 0.00 | 0.00 | 0.17 | 0.17 |
| cold | 90 | 3 | 16.2 | 0.42 | 0.00 | 0.24 | 0.24 |
| cold | 90 | 5 | 12.6 | 0.55 | 0.30 | 0.15 | 0.44 |
| cold | 100 | 1 | 12.1 | 0.18 | 0.10 | 0.20 | 0.30 |
| cold | 100 | 3 | 18.2 | 0.18 | 0.18 | 0.19 | 0.37 |
| cold | 100 | 5 | 13.9 | 0.32 | 0.32 | 0.20 | 0.51 |
| hot | 10 | 1 | 3.9 | 0.00 | 0.00 | 0.08 | 0.08 |
| hot | 10 | 3 | 5.3 | 0.00 | 0.00 | 0.14 | 0.14 |
| hot | 10 | 5 | 4.4 | 0.00 | 0.00 | 0.16 | 0.16 |
| hot | 20 | 1 | 15.7 | 0.00 | 0.00 | 0.30 | 0.30 |
| hot | 20 | 3 | 13.7 | 0.11 | 0.00 | 0.59 | 0.59 |
| hot | 20 | 5 | 15.6 | 0.00 | 0.00 | 0.57 | 0.57 |
| hot | 30 | 1 | 15.9 | 0.22 | 0.09 | 0.45 | 0.53 |
| hot | 30 | 3 | 13.1 | 0.32 | 0.07 | 0.57 | 0.64 |
| hot | 30 | 5 | 16.2 | 0.34 | 0.24 | 0.50 | 0.74 |
| hot | 40 | 1 | 15.7 | 0.00 | 0.00 | 0.33 | 0.33 |
| hot | 40 | 3 | 13.1 | 0.00 | 0.00 | 0.63 | 0.63 |
| hot | 40 | 5 | 16.2 | 0.22 | 0.00 | 0.70 | 0.70 |
| hot | 50 | 1 | 15.7 | 0.00 | 0.00 | 0.32 | 0.32 |
| hot | 50 | 3 | 13.3 | 0.12 | 0.00 | 0.64 | 0.64 |
| hot | 50 | 5 | 15.0 | 0.88 | 0.60 | 0.64 | 1.24 |
| hot | 60 | 1 | 17.4 | 0.27 | 0.00 | 0.44 | 0.44 |
| hot | 60 | 3 | 14.0 | 0.31 | 0.08 | 0.56 | 0.64 |
| hot | 60 | 5 | 15.6 | 0.79 | 0.53 | 0.75 | 1.28 |

Table 6: Coordinates of TAS regions used in this work. We defined the region between the most distal gene and the end of the chromosome as TAS regions. chr. chromosome; spec. species from which the most distal gene (gene) was used, either *D. simulans* or *D. melanogaster*

| name | chr. | start | end | length [bp] | gene | spec. |
|---:|---:|---:|---:|---:|---:|---:|
| X-TAS | X | 1 | 68,388 | 68,388 | GD16489 | Dsim |
| 2L-TAS | 2L | 1 | 193,350 | 193,350 | l(2)gl | Dmel |
| 2R-TAS | 2R | 23,209,012 | 23,239,221 | 30,210 | CG9380 | Dmel |
| 3L-TAS | 3L | 1 | 41,006 | 41,006 | GD13543 | Dsim |
| 3R-TAS | 3R | 28,694,848 | 28,742,805 | 47,958 | Map205 | Dmel |

Table 7: Overview of P-element insertions in TAS regions. For each condition (cond; hot/cold/base), generation (g.) and replicate we show the number of anchor reads mapping to X-TAS, 2L-TAS, 2R-TAS, 3L-TAS and 3R-TAS. Based on an unbiased estimate of the P-element abundance (Pi; using DeviaTE) and the fraction of anchor reads aligning to TAS regions the number of P-element insertions in TAS regions can be computed ($TAS\_insertions = Pi * TAS\_anchor\_reads/an$, where $an$ is the total number of anchor reads). The enrichment of P-element insertions in TAS regions relative to random expectations was computed base on the total length of the genome and the length of the TAS regions.

| | | | | | anchor reads in TAS | | | | | | insertions in TAS | | | | | | enrichment in TAS | | | | | |
|---|---|---|---|---|---|---|---|---|---|---|---|---|---|---|---|---|---|---|---|---|---|---|
| cond | g | r | Pi | an | all | X | 2L | 2R | 3L | 3R | all | X | 2L | 2R | 3L | 3R | all | X | 2L | 2R | 3L | 3R |
| base | 0 | 1 | 0.85 | 251 | 7 | 5 | 1 | 0 | 0 | 1 | 0.02 | 0.02 | 0.00 | 0.00 | 0.00 | 0.00 | 10.7 | 42.8 | 3.0 | 0.0 | 0.0 | 12.2 |
| base | 0 | 3 | 0.80 | 336 | 8 | 4 | 3 | 0 | 0 | 1 | 0.02 | 0.01 | 0.01 | 0.00 | 0.00 | 0.00 | 9.2 | 25.5 | 6.8 | 0.0 | 0.0 | 9.1 |
| base | 0 | 5 | 0.92 | 428 | 11 | 6 | 2 | 0 | 0 | 3 | 0.02 | 0.01 | 0.00 | 0.00 | 0.00 | 0.01 | 9.9 | 30.1 | 3.5 | 0.0 | 0.0 | 21.5 |
| cold | 10 | 1 | 1.21 | 146 | 6 | 2 | 3 | 0 | 0 | 1 | 0.05 | 0.02 | 0.02 | 0.00 | 0.00 | 0.01 | 15.8 | 29.4 | 15.6 | 0.0 | 0.0 | 21.0 |
| cold | 10 | 3 | 1.83 | 230 | 4 | 3 | 1 | 0 | 0 | 0 | 0.03 | 0.02 | 0.01 | 0.00 | 0.00 | 0.00 | 6.7 | 28.0 | 3.3 | 0.0 | 0.0 | 0.0 |
| cold | 10 | 5 | 1.20 | 153 | 10 | 2 | 0 | 0 | 0 | 8 | 0.08 | 0.02 | 0.00 | 0.00 | 0.00 | 0.06 | 25.2 | 28.1 | 0.0 | 0.0 | 0.0 | 160.0 |
| cold | 20 | 1 | 1.87 | 294 | 4 | 1 | 2 | 0 | 0 | 1 | 0.03 | 0.01 | 0.01 | 0.00 | 0.00 | 0.01 | 5.2 | 7.3 | 5.2 | 0.0 | 0.0 | 10.4 |
| cold | 20 | 3 | 2.88 | 270 | 5 | 3 | 1 | 0 | 0 | 1 | 0.05 | 0.03 | 0.01 | 0.00 | 0.00 | 0.01 | 7.1 | 23.8 | 2.8 | 0.0 | 0.0 | 11.3 |
| cold | 20 | 5 | 1.81 | 160 | 1 | 1 | 0 | 0 | 0 | 0 | 0.01 | 0.01 | 0.00 | 0.00 | 0.00 | 0.00 | 2.4 | 13.4 | 0.0 | 0.0 | 0.0 | 0.0 |
| cold | 30 | 1 | 3.17 | 327 | 3 | 2 | 1 | 0 | 0 | 0 | 0.03 | 0.02 | 0.01 | 0.00 | 0.00 | 0.00 | 3.5 | 13.1 | 2.3 | 0.0 | 0.0 | 0.0 |
| cold | 30 | 3 | 5.11 | 538 | 12 | 8 | 4 | 0 | 0 | 0 | 0.11 | 0.08 | 0.04 | 0.00 | 0.00 | 0.00 | 8.6 | 31.9 | 5.6 | 0.0 | 0.0 | 0.0 |
| cold | 30 | 5 | 2.41 | 251 | 13 | 10 | 3 | 0 | 0 | 0 | 0.13 | 0.10 | 0.03 | 0.00 | 0.00 | 0.00 | 20.0 | 85.5 | 9.1 | 0.0 | 0.0 | 0.0 |
| cold | 40 | 1 | 7.89 | 806 | 9 | 9 | 0 | 0 | 0 | 0 | 0.09 | 0.09 | 0.00 | 0.00 | 0.00 | 0.00 | 4.3 | 24.0 | 0.0 | 0.0 | 0.0 | 0.0 |
| cold | 40 | 3 | 12.80 | 3023 | 56 | 31 | 17 | 1 | 0 | 7 | 0.24 | 0.13 | 0.07 | 0.00 | 0.00 | 0.03 | 7.1 | 22.0 | 4.3 | 1.6 | 0.0 | 7.1 |
| cold | 40 | 5 | 5.77 | 609 | 7 | 2 | 2 | 0 | 0 | 3 | 0.07 | 0.02 | 0.02 | 0.00 | 0.00 | 0.03 | 4.4 | 7.0 | 2.5 | 0.0 | 0.0 | 15.1 |
| cold | 50 | 1 | 12.80 | 2007 | 25 | 17 | 6 | 0 | 0 | 2 | 0.16 | 0.11 | 0.04 | 0.00 | 0.00 | 0.01 | 4.8 | 18.2 | 2.3 | 0.0 | 0.0 | 3.0 |
| cold | 50 | 3 | 16.78 | 2862 | 37 | 19 | 12 | 0 | 0 | 6 | 0.22 | 0.11 | 0.07 | 0.00 | 0.00 | 0.04 | 5.0 | 14.2 | 3.2 | 0.0 | 0.0 | 6.4 |
| cold | 50 | 5 | 8.54 | 1136 | 17 | 10 | 5 | 0 | 0 | 2 | 0.13 | 0.08 | 0.04 | 0.00 | 0.00 | 0.02 | 5.8 | 18.9 | 3.3 | 0.0 | 0.0 | 5.4 |
| cold | 60 | 1 | 15.16 | 914 | 27 | 15 | 7 | 0 | 0 | 5 | 0.45 | 0.25 | 0.12 | 0.00 | 0.00 | 0.08 | 11.4 | 35.2 | 5.8 | 0.0 | 0.0 | 16.7 |
| cold | 60 | 3 | 22.46 | 1594 | 19 | 6 | 7 | 0 | 0 | 6 | 0.27 | 0.08 | 0.10 | 0.00 | 0.00 | 0.08 | 4.6 | 8.1 | 3.3 | 0.0 | 0.0 | 11.5 |
| cold | 60 | 5 | 13.55 | 2437 | 19 | 8 | 8 | 0 | 0 | 3 | 0.11 | 0.04 | 0.04 | 0.00 | 0.00 | 0.02 | 3.0 | 7.0 | 2.5 | 0.0 | 0.0 | 3.8 |
| cold | 70 | 1 | 15.25 | 2091 | 44 | 22 | 18 | 0 | 0 | 4 | 0.32 | 0.16 | 0.13 | 0.00 | 0.00 | 0.03 | 8.1 | 22.6 | 6.5 | 0.0 | 0.0 | 5.9 |
| cold | 70 | 3 | 21.65 | 4469 | 75 | 44 | 23 | 0 | 0 | 8 | 0.36 | 0.21 | 0.11 | 0.00 | 0.00 | 0.04 | 6.5 | 21.1 | 3.9 | 0.0 | 0.0 | 5.5 |
| cold | 70 | 5 | 15.49 | 2700 | 40 | 18 | 7 | 0 | 0 | 15 | 0.23 | 0.10 | 0.04 | 0.00 | 0.00 | 0.09 | 5.7 | 14.3 | 2.0 | 0.0 | 0.0 | 17.0 |
| cold | 80 | 1 | 12.77 | 2015 | 35 | 15 | 19 | 0 | 0 | 1 | 0.22 | 0.10 | 0.12 | 0.00 | 0.00 | 0.01 | 6.7 | 16.0 | 7.2 | 0.0 | 0.0 | 1.5 |
| cold | 80 | 3 | 18.08 | 3066 | 38 | 30 | 7 | 0 | 0 | 1 | 0.22 | 0.18 | 0.04 | 0.00 | 0.00 | 0.01 | 4.8 | 21.0 | 1.7 | 0.0 | 0.0 | 1.0 |
| cold | 80 | 5 | 13.63 | 2223 | 44 | 17 | 11 | 0 | 0 | 16 | 0.27 | 0.10 | 0.07 | 0.00 | 0.00 | 0.10 | 7.6 | 16.4 | 3.8 | 0.0 | 0.0 | 22.0 |
| cold | 90 | 1 | 11.93 | 3672 | 53 | 20 | 17 | 0 | 0 | 16 | 0.17 | 0.06 | 0.06 | 0.00 | 0.00 | 0.05 | 5.6 | 11.7 | 3.5 | 0.0 | 0.0 | 13.3 |
| cold | 90 | 3 | 16.24 | 4416 | 64 | 47 | 8 | 0 | 0 | 9 | 0.24 | 0.17 | 0.03 | 0.00 | 0.00 | 0.03 | 5.6 | 22.8 | 1.4 | 0.0 | 0.0 | 6.2 |
| cold | 90 | 5 | 12.62 | 2465 | 29 | 11 | 5 | 0 | 0 | 13 | 0.15 | 0.06 | 0.03 | 0.00 | 0.00 | 0.07 | 4.5 | 9.6 | 1.5 | 0.0 | 0.0 | 16.1 |
| cold | 100 | 1 | 12.14 | 2929 | 48 | 27 | 4 | 0 | 0 | 17 | 0.20 | 0.11 | 0.02 | 0.00 | 0.00 | 0.07 | 6.3 | 19.8 | 1.0 | 0.0 | 0.0 | 17.8 |
| cold | 100 | 3 | 18.16 | 4400 | 46 | 33 | 10 | 0 | 0 | 3 | 0.19 | 0.14 | 0.04 | 0.00 | 0.00 | 0.01 | 4.0 | 16.1 | 1.7 | 0.0 | 0.0 | 2.1 |
| cold | 100 | 5 | 13.87 | 3037 | 43 | 13 | 17 | 0 | 0 | 13 | 0.20 | 0.06 | 0.08 | 0.00 | 0.00 | 0.06 | 5.5 | 9.2 | 4.2 | 0.0 | 0.0 | 13.1 |
| hot | 10 | 1 | 3.89 | 503 | 10 | 5 | 4 | 0 | 0 | 1 | 0.08 | 0.04 | 0.03 | 0.00 | 0.00 | 0.01 | 7.7 | 21.3 | 6.0 | 0.0 | 0.0 | 6.1 |
| hot | 10 | 3 | 5.32 | 514 | 14 | 11 | 2 | 0 | 0 | 1 | 0.14 | 0.11 | 0.02 | 0.00 | 0.00 | 0.01 | 10.5 | 45.9 | 3.0 | 0.0 | 0.0 | 6.0 |
| hot | 10 | 5 | 4.38 | 779 | 29 | 26 | 1 | 1 | 0 | 1 | 0.16 | 0.15 | 0.01 | 0.01 | 0.00 | 0.01 | 14.3 | 71.6 | 1.0 | 6.2 | 0.0 | 3.9 |
| hot | 20 | 1 | 15.73 | 4333 | 82 | 53 | 20 | 0 | 0 | 9 | 0.30 | 0.19 | 0.07 | 0.00 | 0.00 | 0.03 | 7.3 | 26.3 | 3.5 | 0.0 | 0.0 | 6.4 |
| hot | 20 | 3 | 13.68 | 3669 | 158 | 90 | 33 | 0 | 0 | 35 | 0.59 | 0.34 | 0.12 | 0.00 | 0.00 | 0.13 | 16.6 | 52.6 | 6.8 | 0.0 | 0.0 | 29.2 |
| hot | 20 | 5 | 15.59 | 4354 | 158 | 91 | 54 | 0 | 1 | 12 | 0.57 | 0.33 | 0.19 | 0.00 | 0.00 | 0.04 | 14.0 | 44.9 | 9.4 | 0.0 | 0.8 | 8.4 |
| hot | 30 | 1 | 15.94 | 2866 | 80 | 55 | 12 | 0 | 0 | 13 | 0.45 | 0.31 | 0.07 | 0.00 | 0.00 | 0.07 | 10.8 | 41.2 | 3.2 | 0.0 | 0.0 | 13.9 |
| hot | 30 | 3 | 13.08 | 2124 | 93 | 57 | 15 | 0 | 0 | 21 | 0.57 | 0.35 | 0.09 | 0.00 | 0.00 | 0.13 | 16.9 | 57.6 | 5.4 | 0.0 | 0.0 | 30.3 |
| hot | 30 | 5 | 16.20 | 1646 | 51 | 36 | 11 | 0 | 0 | 4 | 0.50 | 0.35 | 0.11 | 0.00 | 0.00 | 0.04 | 11.9 | 46.9 | 5.1 | 0.0 | 0.0 | 7.4 |
| hot | 40 | 1 | 15.69 | 3276 | 68 | 36 | 23 | 0 | 0 | 9 | 0.33 | 0.17 | 0.11 | 0.00 | 0.00 | 0.04 | 8.0 | 23.6 | 5.3 | 0.0 | 0.0 | 8.4 |
| hot | 40 | 3 | 13.09 | 3697 | 178 | 87 | 52 | 0 | 3 | 36 | 0.63 | 0.31 | 0.18 | 0.00 | 0.01 | 0.13 | 18.6 | 50.5 | 10.7 | 0.0 | 2.9 | 29.8 |
| hot | 40 | 5 | 16.24 | 2636 | 114 | 59 | 34 | 0 | 0 | 21 | 0.70 | 0.36 | 0.21 | 0.00 | 0.00 | 0.13 | 16.7 | 48.0 | 9.8 | 0.0 | 0.0 | 24.4 |
| hot | 50 | 1 | 15.74 | 1575 | 32 | 15 | 8 | 0 | 0 | 9 | 0.32 | 0.15 | 0.08 | 0.00 | 0.00 | 0.09 | 7.8 | 20.4 | 3.9 | 0.0 | 0.0 | 17.5 |
| hot | 50 | 3 | 13.32 | 1324 | 64 | 35 | 18 | 0 | 0 | 11 | 0.64 | 0.35 | 0.18 | 0.00 | 0.00 | 0.11 | 18.6 | 56.7 | 10.3 | 0.0 | 0.0 | 25.4 |
| hot | 50 | 5 | 15.04 | 1744 | 74 | 32 | 32 | 0 | 1 | 9 | 0.64 | 0.28 | 0.28 | 0.00 | 0.01 | 0.08 | 16.3 | 39.4 | 13.9 | 0.0 | 2.1 | 15.8 |
| hot | 60 | 1 | 17.35 | 3058 | 78 | 36 | 13 | 0 | 0 | 29 | 0.44 | 0.20 | 0.07 | 0.00 | 0.00 | 0.16 | 9.8 | 25.3 | 3.2 | 0.0 | 0.0 | 29.0 |
| hot | 60 | 3 | 14.04 | 1541 | 62 | 29 | 20 | 0 | 1 | 12 | 0.56 | 0.26 | 0.18 | 0.00 | 0.01 | 0.11 | 15.5 | 40.4 | 9.9 | 0.0 | 2.3 | 23.8 |
| hot | 60 | 5 | 15.64 | 1722 | 83 | 42 | 31 | 0 | 0 | 10 | 0.75 | 0.38 | 0.28 | 0.00 | 0.00 | 0.09 | 18.6 | 52.3 | 13.7 | 0.0 | 0.0 | 17.8 |

Table 8: Upper estimate of the number of piRNA-producing P-element insertions (sum) in the experimental populations. Anchor reads were mapped to the sequences of TAS regions and piRNA clusters. We used both ambiguously and unambiguously mapped anchor reads. Also the annotation of the piRNA clusters was based on ambiguously and unambiguously aligned piRNAs.

| | | | | anchor | | | insertions | | |
|---|---|---|---|---|---|---|---|---|---|
| | g | r | Pi | all | clu | tas | clu | tas | sum |
| base | 0 | 1 | 0.85 | 251 | 11 | 6 | 0.04 | 0.02 | 0.06 |
| base | 0 | 3 | 0.80 | 336 | 22 | 8 | 0.05 | 0.02 | 0.07 |
| base | 0 | 5 | 0.92 | 427 | 31 | 7 | 0.07 | 0.02 | 0.08 |
| cold | 10 | 1 | 1.21 | 146 | 14 | 4 | 0.12 | 0.03 | 0.15 |
| cold | 10 | 3 | 1.83 | 230 | 16 | 2 | 0.13 | 0.02 | 0.14 |
| cold | 10 | 5 | 1.20 | 153 | 8 | 9 | 0.06 | 0.07 | 0.13 |
| cold | 20 | 1 | 1.87 | 295 | 14 | 4 | 0.09 | 0.03 | 0.11 |
| cold | 20 | 3 | 2.88 | 270 | 17 | 4 | 0.18 | 0.04 | 0.22 |
| cold | 20 | 5 | 1.81 | 160 | 12 | 1 | 0.14 | 0.01 | 0.15 |
| cold | 30 | 1 | 3.17 | 327 | 31 | 1 | 0.30 | 0.01 | 0.31 |
| cold | 30 | 3 | 5.11 | 538 | 35 | 7 | 0.33 | 0.07 | 0.40 |
| cold | 30 | 5 | 2.41 | 248 | 19 | 6 | 0.18 | 0.06 | 0.24 |
| cold | 40 | 1 | 7.89 | 806 | 35 | 5 | 0.34 | 0.05 | 0.39 |
| cold | 40 | 3 | 12.80 | 3023 | 193 | 31 | 0.82 | 0.13 | 0.95 |
| cold | 40 | 5 | 5.77 | 610 | 60 | 5 | 0.57 | 0.05 | 0.61 |
| cold | 50 | 1 | 12.80 | 2008 | 117 | 21 | 0.75 | 0.13 | 0.88 |
| cold | 50 | 3 | 16.78 | 2867 | 182 | 26 | 1.07 | 0.15 | 1.22 |
| cold | 50 | 5 | 8.54 | 1136 | 90 | 9 | 0.68 | 0.07 | 0.74 |
| cold | 60 | 1 | 15.16 | 914 | 88 | 21 | 1.46 | 0.35 | 1.81 |
| cold | 60 | 3 | 22.46 | 1594 | 191 | 8 | 2.69 | 0.11 | 2.80 |
| cold | 60 | 5 | 13.55 | 2443 | 268 | 9 | 1.49 | 0.05 | 1.54 |
| cold | 70 | 1 | 15.25 | 2091 | 123 | 35 | 0.90 | 0.26 | 1.15 |
| cold | 70 | 3 | 21.65 | 4479 | 441 | 46 | 2.13 | 0.22 | 2.35 |
| cold | 70 | 5 | 15.49 | 2705 | 209 | 27 | 1.20 | 0.15 | 1.35 |
| cold | 80 | 1 | 12.77 | 2030 | 138 | 33 | 0.87 | 0.21 | 1.08 |
| cold | 80 | 3 | 18.08 | 3088 | 297 | 21 | 1.74 | 0.12 | 1.86 |
| cold | 80 | 5 | 13.63 | 2234 | 160 | 36 | 0.98 | 0.22 | 1.20 |
| cold | 90 | 1 | 11.93 | 3678 | 354 | 39 | 1.15 | 0.13 | 1.27 |
| cold | 90 | 3 | 16.24 | 4421 | 392 | 40 | 1.44 | 0.15 | 1.59 |
| cold | 90 | 5 | 12.62 | 2471 | 262 | 18 | 1.34 | 0.09 | 1.43 |
| cold | 100 | 1 | 12.14 | 2935 | 219 | 36 | 0.91 | 0.15 | 1.05 |
| cold | 100 | 3 | 18.16 | 4414 | 379 | 32 | 1.56 | 0.13 | 1.69 |
| cold | 100 | 5 | 13.87 | 3042 | 323 | 26 | 1.47 | 0.12 | 1.59 |
| hot | 10 | 1 | 3.89 | 504 | 38 | 5 | 0.29 | 0.04 | 0.33 |
| hot | 10 | 3 | 5.32 | 511 | 42 | 4 | 0.44 | 0.04 | 0.48 |
| hot | 10 | 5 | 4.38 | 768 | 58 | 6 | 0.33 | 0.03 | 0.36 |
| hot | 20 | 1 | 15.73 | 4326 | 338 | 52 | 1.23 | 0.19 | 1.42 |
| hot | 20 | 3 | 13.68 | 3645 | 233 | 96 | 0.87 | 0.36 | 1.23 |
| hot | 20 | 5 | 15.59 | 4340 | 302 | 109 | 1.08 | 0.39 | 1.48 |
| hot | 30 | 1 | 15.94 | 2848 | 262 | 39 | 1.47 | 0.22 | 1.69 |
| hot | 30 | 3 | 13.08 | 2110 | 193 | 49 | 1.20 | 0.30 | 1.50 |
| hot | 30 | 5 | 16.20 | 1643 | 191 | 34 | 1.88 | 0.34 | 2.22 |
| hot | 40 | 1 | 15.69 | 3270 | 242 | 43 | 1.16 | 0.21 | 1.37 |
| hot | 40 | 3 | 13.09 | 3691 | 291 | 153 | 1.03 | 0.54 | 1.57 |
| hot | 40 | 5 | 16.24 | 2630 | 209 | 88 | 1.29 | 0.54 | 1.83 |
| hot | 50 | 1 | 15.74 | 1574 | 127 | 25 | 1.27 | 0.25 | 1.52 |
| hot | 50 | 3 | 13.32 | 1321 | 91 | 52 | 0.92 | 0.52 | 1.44 |
| hot | 50 | 5 | 15.04 | 1744 | 158 | 59 | 1.36 | 0.51 | 1.87 |
| hot | 60 | 1 | 17.35 | 3052 | 182 | 59 | 1.03 | 0.34 | 1.37 |
| hot | 60 | 3 | 14.04 | 1541 | 140 | 53 | 1.28 | 0.48 | 1.76 |
| hot | 60 | 5 | 15.64 | 1724 | 163 | 71 | 1.48 | 0.64 | 2.12 |

Table 9: Fraction of anchor reads that could not be aligned to the reference genome (in %). cond condition, g generation

| | | replicate | | |
|---|---|---|---|---|
| cond | g | 1 | 3 | 5 |
| base | 0 | 6.0 | 11.6 | 9.4 |
| cold | 10 | 8.9 | 11.7 | 11.1 |
| cold | 20 | 10.9 | 7.4 | 8.8 |
| cold | 30 | 11.9 | 10.4 | 10.1 |
| cold | 40 | 6.6 | 8.4 | 6.2 |
| cold | 50 | 6.3 | 5.8 | 7.6 |
| cold | 60 | 9.1 | 7.2 | 8.6 |
| cold | 70 | 5.3 | 6.8 | 7.0 |
| cold | 80 | 5.7 | 5.3 | 7.1 |
| cold | 90 | 6.9 | 7.2 | 6.6 |
| cold | 100 | 7.9 | 8.5 | 10.5 |
| hot | 10 | 8.9 | 9.2 | 12.1 |
| hot | 20 | 11.3 | 12.0 | 9.9 |
| hot | 30 | 17.1 | 17.3 | 12.1 |
| hot | 40 | 9.5 | 10.2 | 10.8 |
| hot | 50 | 7.4 | 7.5 | 8.2 |
| hot | 60 | 12.8 | 11.5 | 13.1 |
